# Comprehensive evaluation of phosphoproteomic-based kinase activity inference

**DOI:** 10.1101/2024.06.27.601117

**Authors:** Sophia Müller-Dott, Eric J. Jaehnig, Khoi Pham Munchic, Wen Jiang, Tomer M. Yaron-Barir, Sara R. Savage, Martin Garrido-Rodriguez, Jared L. Johnson, Alessandro Lussana, Evangelia Petsalaki, Jonathan T. Lei, Aurelien Dugourd, Karsten Krug, Lewis C. Cantley, D.R. Mani, Bing Zhang, Julio Saez-Rodriguez

## Abstract

Kinases play a central role in regulating cellular processes, making their study essential for understanding cellular function and disease mechanisms. To investigate the regulatory state of a kinase, numerous methods have been, and continue to be, developed to infer kinase activities from phosphoproteomics data. These methods usually rely on a set of kinase targets collected from various kinase-substrate libraries. However, only a small percentage of measured phosphorylation sites can usually be attributed to an upstream kinase in these libraries, limiting the scope of kinase activity inference. In addition, the inferred activities from different methods can vary making it crucial to evaluate them for accurate interpretation. Here, we present a comprehensive evaluation of kinase activity inference methods using multiple kinase-substrate libraries combined with different inference algorithms. Additionally, we try to overcome the coverage limitations for measured targets in kinase substrate libraries by adding predicted kinase-substrate interactions for activity inference. For the evaluation, in addition to classical cell-based perturbation experiments, we introduce a tumor-based benchmarking approach that utilizes multi-omics data to identify highly active or inactive kinases per tumor type. We show that while most computational algorithms perform comparably regardless of their complexity, the choice of kinase-substrate library can highly impact the inferred kinase activities. Hereby, manually curated libraries, particularly PhosphoSitePlus, demonstrate superior performance in recapitulating kinase activities from phosphoproteomics data. Additionally, in the tumor-based evaluation, adding predicted targets from NetworKIN further boosts the performance, while normalizing sites to host protein levels reduces kinase activity inference performance. We then showcase how kinase activity inference can help in characterizing the response to kinase inhibitors in different cell lines. Overall, the selection of reliable kinase activity inference methods is important in identifying deregulated kinases and novel drug targets. Finally, to facilitate the evaluation of novel methods in the future, we provide both benchmarking approaches in the R package benchmarKIN.

**Graphical Abstract:** 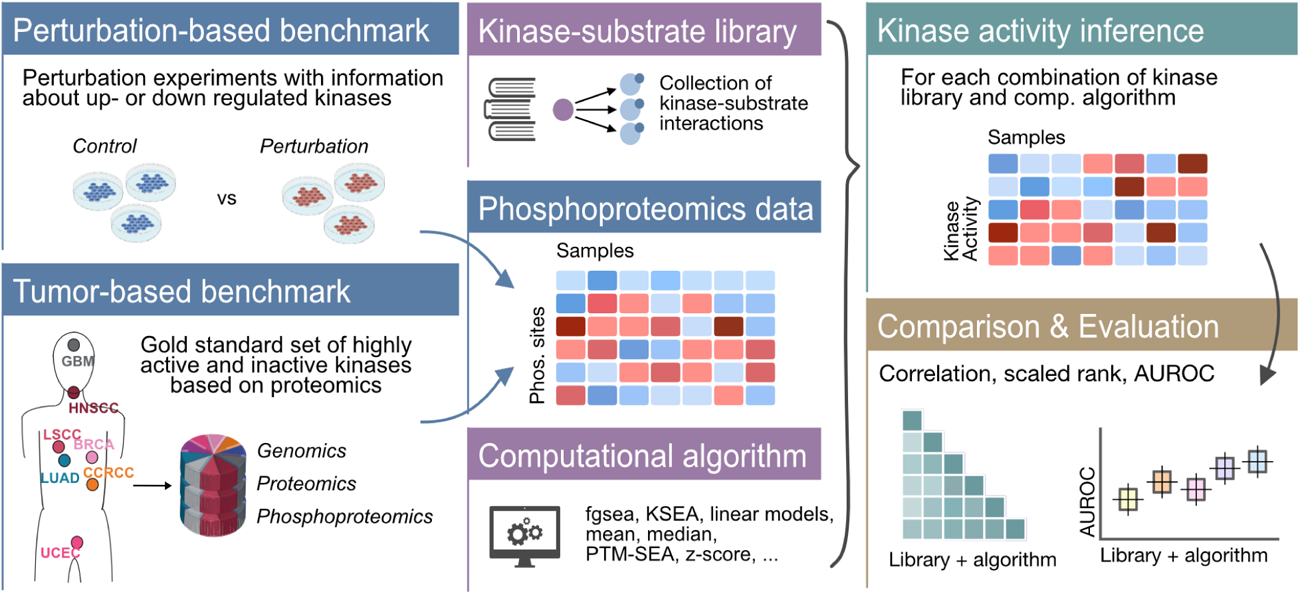

## Introduction

Protein phosphorylation is a reversible post-translational modification that acts as a key regulator of various cellular processes and plays a central role in intracellular signal transduction^1^. It is controlled by kinases which, together with their substrates and phosphatases, form a large network that controls diverse biological processes ranging from cell cycle progression, cell growth and differentiation to apoptosis. There are roughly 540 kinases encoded in the human genome that phosphorylate 20,000 proteins at more than 350,000 phosphorylation sites^2^. By catalyzing the transfer of a phosphate group to threonine, serine, tyrosine or histidine residues, they affect the substrate protein’s activity, stability, localization and/or interaction with other molecules^3^. Aberrant kinase activity has been implicated in the pathogenesis of numerous diseases, including Alzheimer’s disease^4^, Parkinson’s disease^5^, metabolic dysfunction-associated steatotic liver disease^6^, obesity and diabetes^7^, as well as various cancer types^8^. Protein kinases are also one of the most targeted protein families for inhibition by small molecules^9^. Hence, investigating the regulatory state of a kinase has emerged as an important objective in many biomedical contexts, including identification of novel disease-specific drug targets, development of patient specific therapeutics and prediction of treatment outcomes^10–12^.

Enabled by mass spectrometry (MS)-based technologies, measuring global phosphorylation events has provided new opportunities for the systematic analysis of kinases and their activities. Large-scale identification and quantification of phosphorylation levels can be obtained by MS, which can provide measurements for up to 50,000 unique phospho-peptides that span over 75% of all cellular proteins^13^. This snapshot of the phosphoproteome reflects the activity of kinases and phosphatases. For example, to better understand dysregulation of phosphorylation in cancer, phosphoproteomic profiling approaches have been routinely applied to tumor cohorts in Clinical Proteogenomic Tumor Analysis Consortium (CPTAC)^14,15^ and International Cancer Proteogenome Consortium (ICPC)^16^ studies. Additionally, phosphoproteomic profiling has recently been applied to better understand the effects of a SARS-CoV-2 infection on cellular signaling^17^.

Phosphoproteomics data can be used to infer the activity of a given kinase based on the phosphorylation profiles of the kinase’s targets^18^. Several tools have been developed to carry out this inference utilizing computational algorithms with varying complexity. For example, PTM-SEA^19^, uses the single-sample gene set enrichment algorithm^20^, whereas KSEA^21,22^ calculates a z-score based on the aggregation of phosphorylation site levels for known targets relative to the background set of overall phosphorylation site levels. A common feature of all these methods is that prior knowledge of the target phosphorylation sites of the respective kinase is required. Typically, the kinase target site sets are extracted from manually curated databases of known targets such as PhosphoSitePlus^2^, SIGNOR^23^, or Phospho.ELM^24^. However, only a small percentage of phosphorylation sites can be attributed to any kinase, and, of those that can be, many are often attributed to a small handful of well-studied kinases^25,26^. This can affect kinase activity inference since activity cannot be inferred if too few substrates of a given kinase are measured, and inferences that are based on low numbers of substrates may not be as accurate as those made using a greater number of reliable substrates. Therefore, it is critical to determine if we can increase the number of assessable kinases and enhance performance by boosting the number of potential substrates that are considered. One way to accomplish this is by including sites identified as targets in large *in vitro* screening assays^27,28^. Alternatively, one could consider sites predicted as targets by computational tools, such as NetworKIN^29^.

Given that multiple different methods have been, and continue to be, developed to infer kinase activity and that the performance of these methods is dependent on the kinase targets sets that are chosen, it is critical to establish mechanisms to evaluate the kinase activity inference performance in order to determine the optimal approaches for estimating kinase activity from phosphoproteomics data. So far, smaller comparative analyses have been performed that relied on perturbation studies aimed at identifying perturbed kinases from phosphoproteomic data^30,31^. However, using perturbation studies alone for evaluation has been limited to a subset of well-studied kinases, are only available in an *in vitro* setting, and may be affected by unknown off-target effects.

In this study, we present two complementary benchmarking strategies designed to evaluate various combinations of computational algorithms and kinase-substrate libraries for inferring kinase activities. For the kinase activity inference, we focused on methods that calculate a score solely based on kinase-substrate interactions, can be applied for single-sample analysis as well as comparative analysis and do not require a specific input file. As such we have not considered methods, such as CLUE^32^, KinasePA^33^, KEA3^34^, INKA^35^ or KSTAR^36^ from the comparison. For the cell-based kinase perturbation-based evaluation, we expand the existing gold-standard benchmark set of perturbation experiments to encompass a broader set of kinases for evaluation. Furthermore, we introduce a new tumor-based benchmarking approach based on the multi-omics CPTAC datasets to ultimately determine the optimal methodology for inferring kinase activities in human tumors. We implement these benchmarking approaches in the R package benchmarKIN (https://github.com/saezlab/benchmarKIN) to facilitate the evaluation of novel methodologies.

## Results

### Quantifying kinome coverage by prior knowledge resources

For the comparison of kinase activity estimation we collected six kinase-substrate libraries: PhosphoSitePlus^2^, PTMsigDB^19^, the gold standard set used to train GPS 5.0 (GPS gold)^37^, OmniPath^38^, iKiP-DB^27^ and NetworKIN^29^. Besides manually curated resources and meta-resources, like PhosphoSitePlus, PTMsigDB, GPS gold and OmniPath, we also included the *in vitro* database iKiP-DB (*in vitro* Kinase-to-Phosphosite database), which identified kinase-substrate interactions from a large-scale *in vitro* kinase study for over 300 human proteins, and kinase target sites from the NetworKIN database, which contains precomputed kinase-substrate interactions for phosphorylation sites reported in the KinomeXplorer-DB^39^ using the NetworKIN algorithm^29^.

We then compared the coverage of kinases and kinase-substrate interactions for the different libraries. OmniPath, which is a meta-resource that includes interactions from dbPTM^40^, HPRD^41^, Li2012^42^, MIMP^43^, NCI-PID^44^, PhosphoNetworks^45^, PhosphoSitePlus^2^, phospho.ELM^24^, Reactome^46^, RLIMS-P^47^, and SIGNOR^23^, exhibited the highest kinase coverage (467), followed by PhosphoSitePlus (390) and GPS gold (352) (Fig. 1a). OmniPath includes 47 kinases not present in any of the other resources that mainly originate from kinases with interactions reported by MIMP and PhosphoNetworks. iKiP-DB, PhosphoSitePlus, NetworKIN and GPS gold also report interactions for 11, 7, 2 and 1 unique kinases, respectively. 86.2% of all kinases are covered by at least two of the analyzed resources (Fig. 1a, Supplementary Fig. 1a). In general, all databases cover both serine/threonine and tyrosine kinases as well as kinases with ambiguous amino acid specificity (Supplementary Fig. 1b). In terms of kinase-substrate interactions, iKiP-DB and NetworKIN had among the highest number of interactions (iKiP-DB: 26,786, NetworKIN: 22,788), which is expected since interactions from these resources are not limited to those that are experimentally validated. Additionally, the meta-resource OmniPath contained a total of 26,280 interactions (Fig. 1b). A lower overlap of kinase-substrate interactions than of kinase coverage was observed between the resources, with only 21.7% of interactions being shared between at least two resources. iKiP-DB and NetworKIN had the lowest overlap with the other resources and reported 26,327 and 19,524 unique kinase-substrate interactions, respectively. OmniPath, PhosphoSitePlus, PTMsigDB and GPS gold contained 11,148, 341, 277 and 544 unique kinase-substrate interactions, respectively (Fig. 1b, Supplementary Fig. 1c). In general, the manually curated resources, PhosphoSitePlus, PTMsigDB, GPS gold and OmniPath all have a median number of targets between 8.5 and 18 for all kinases. In contrast, NetworKIN and iKiP-DB, have much larger median numbers of predicted targets across kinases, 64 and 69, respectively (Supplementary Fig. 1d). Lastly, we compared the overlap of targets for each kinase between the resources by calculating the mean Jaccard index of all shared kinases between two resources. We observed higher Jaccard indices between the curated resources, namely PTMsigDB, GPS gold and PhosphoSitePlus. This can be linked to the fact that both PTMsigDB and GPS gold incorporate sites from PhosphoSitePlus. Additionally, with a mean Jaccard index of 0.34, OmniPath shows some overlap with PTMsigDB, GPS gold and PhosphoSitePlus. iKiP-DB and NetworKIN, on the other hand, showed low overlap with any other resource with a highest mean Jaccard index of 0.03 (Fig. 1c).

Overall, we observed variation in kinase coverage among the different resources, with OmniPath exhibiting the most comprehensive kinase coverage, while iKiP-DB had the highest number of kinase-substrate interactions. Additionally, while manually curated databases had overlapping substrate sets, OnmiPath, NetworKIN, and iKiP-DB had substantial numbers of unique substrates, a factor that is likely to impact the accuracy of predicted kinase activities derived from each resource and thus motivates their comparison.

**Figure 1.**
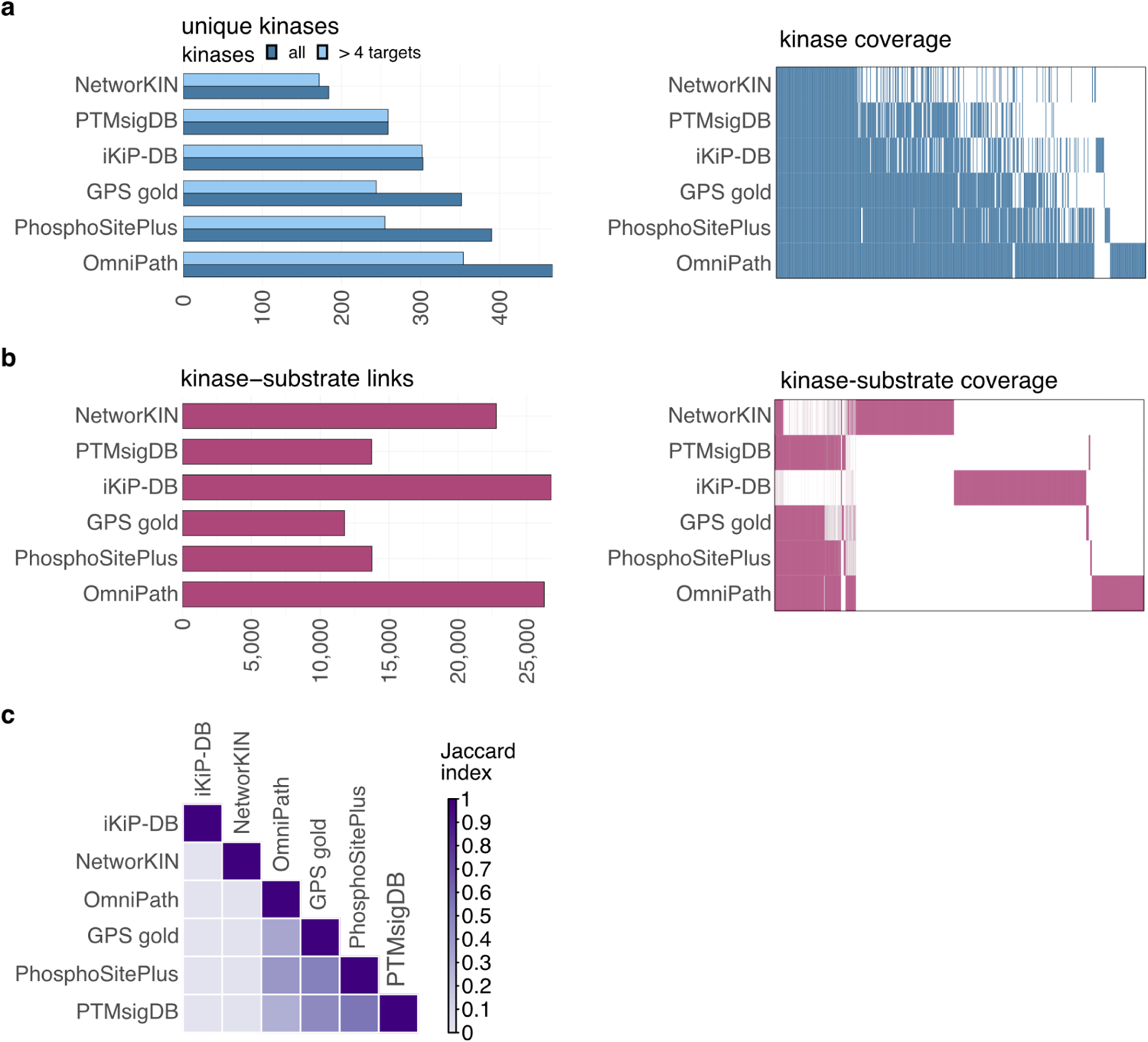
Comparison of kinase-substrate libraries. **a** Number of unique kinases and kinases with at least five annotated targets in each library (left). Coverage of kinases across different kinase-substrate libraries (right). **b** Number of unique kinase-substrate interactions in each library (left) and coverage of kinase-substrate interactions across libraries (right). **c** Mean Jaccard index of kinase regulons between kinase-substrate libraries. For all shared kinases between two libraries, Jaccard indices of their targets were calculated and averaged.

### Unification and correlation analysis of methods for kinase activity inference

To better understand how the different resources may influence kinase activity inference, we used them systematically as alternative input to various computational methods to infer kinase activities from phosphoproteomics datasets.

For the activity estimation, we assessed 17 different methods that predict kinase activity scores from phosphoproteomics data relying on a set of kinase-substrate interactions, namely fgsea^48^ (fast gene set enrichment analysis), KARP^49^ (Kinase activity ranking using phosphoproteomics data), KSEA^21,22^ (Kinase-Substrate Enrichment Analysis), the Kologomorov-Smirnov test (KS test), the linear model implemented in RoKAI^31^ (lm RoKAI), the Mann-Whitney-U test (MWU test), the mean, the median, a multivariate linear model^50^ (mlm), the normalized mean (norm mean), PCA (principal component analysis), PTM-SEA^19^ (PTM-Signature Enrichment Analysis), the sum, a univariate linear model^50^ (ulm), the upper quantile (UQ), VIPER^51^ (Virtual Inference of Protein-activity by Enriched Regulon analysis) and the z-score as implemented by RoKAI^31^ (z-score) (Table 1). Among other things, these methods vary in whether they consider quantitative information, model kinase promiscuity, meaning whether they consider that sites can be phosphorylated by multiple kinases, or whether they calculate a score based on multiple samples. Furthermore, they can be divided into methods that aggregate values for the target sites of a given kinase or compare them to the remaining sites or to an empirical null distribution. A more detailed description of each computational method can be found in the methods section *“Computational methods for kinase activity inference”*.

**Table 1.** Overview of computational methods for kinase activity inference.

| Method | Accounts for magnitude | Models kinase promiscuity | Multi-sample based | Scores relative to |
| --- | --- | --- | --- | --- |
| fgsea<sup>48</sup> | Yes | No | No | Non-targets |
| KARP<sup>49</sup> | Yes | No | No | Non-targets |
| KSEA<sup>21,22</sup> | Yes | No | No | Non-targets |
| Kologomorov-Smirnov<sup>69</sup> | No | No | No | Non-targets |
| Linear model - RoKAI<sup>31</sup> | Yes | Yes | No | Non-targets |
| Mann-Whitney-U | No | No | No | Non-targets |
| mean | Yes | No | No | Solely target based |
| median | Yes | No | No | Solely target based |
| multivariate linear model<sup>50</sup> | Yes | Yes | No | Non-targets |
| normalized mean<sup>50</sup> | Yes | No | No | Null distribution |
| Principal component analysis | Yes | No | Yes | Solely target based |
| PTM-SEA<sup>19</sup> | Yes | No | No | Non-targets & Null distribution |
| sum | Yes | No | No | Solely target based |
| univariate linear model<sup>50</sup> | Yes | No | No | Non-targets |
| upper quantile | Yes | No | No | Solely target based |
| VIPER<sup>51</sup> | Yes | No | No | Non-targets & Null distribution |
| z-score<sup>31</sup> | Yes | No | No | Non-targets |

We then employed each computational method in conjunction with each of the resources described in the previous section on the datasets collected by Hernandez-Armenta et al.^30^ and the datasets presented by Hijazi et al.^52^ (Fig. 2a). This combined collection consists of 212 kinase perturbation experiments with an average of 6,470 phosphorylation sites identified per experiment. Among these, an average of 27% could be assigned to an upstream regulatory kinase in at least one of the kinase-substrate libraries tested (Supplementary Fig. 2a). We then compared the inferred activity scores across experiments when using different computational methods and prior knowledge resources by evaluating mean Pearson correlation coefficients between the different calculation methods and kinase substrate libraries.

Among the computational methods, most showed strong agreement, with a correlation above 0.78 in 80% of cases. The lowest concordance was observed for activity scores inferred using the KARP score, with correlations ranging from 0.03 to -0.19 compared to other methods (Fig. 2b). For the kinase-substrate libraries, we found the highest correlation between PTMsigDB, GPS gold, and PhosphoSitePlus (greater than 0.83). Activity scores inferred with OmniPath also showed a high correlation of over 0.72 with these libraries. However, NetworKIN and iKiP-DB exhibited Pearson correlations below 0.43 when compared to any of the other kinase-substrate libraries (Fig. 2c), which may be expected given that the poor overlap between substrates from these databases and the other databases. In summary, we observed that while most computational algorithms perform comparably, a high discrepancy can be observed between inferred activity scores when using different kinase-substrate libraries.

**Figure 2.**
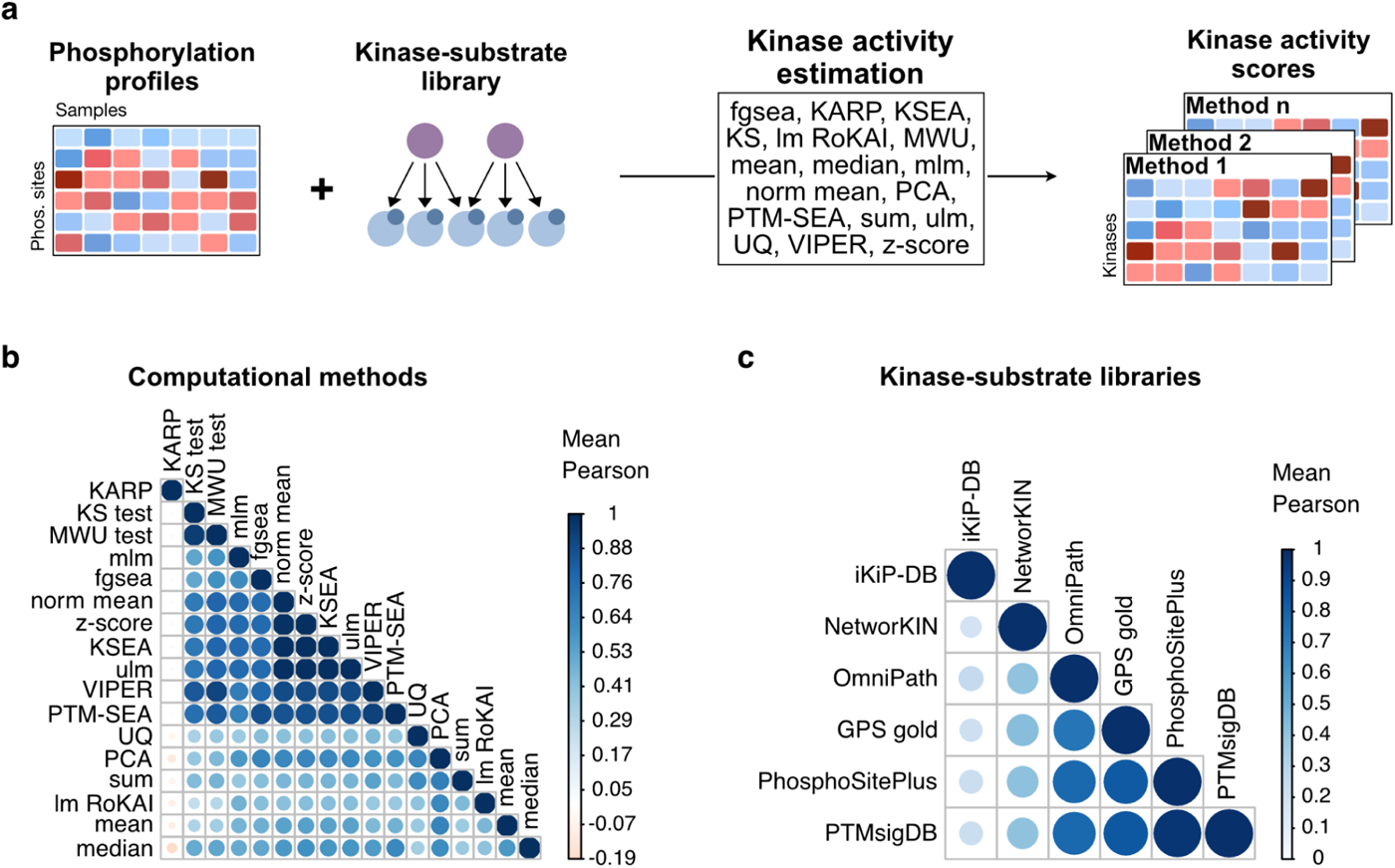
Kinase activity inference comparison. **a** Workflow for kinase activity inference. Using each kinase-substrate library, kinase activity scores are inferred from phosphorylation profiles of 212 different experiments using the following computational methods: fgsea, KARP, KSEA, KS, lm RoKAI, MWU, mean, median, mlm, norm mean, PCA, PTM-SEA, sum, ulm, UQ, VIPER, z-score. **b** Mean Pearson correlation of inferred activity scores. Pearson correlation between computational methods was calculated for each kinase-substrate library and averaged across libraries. **c** Mean Pearson correlation of inferred activity scores. Pearson correlation between kinase-substrate libraries was calculated for each computational method and averaged across methods.

### Benchmarking kinase activity estimation via phosphoproteomics data from experimental perturbations

Next, we evaluated how well each method-resource combination for inferring kinase activity performed with respect to its specific ability to recapitulate the changes in phosphorylation site abundance caused by experimental perturbation of a kinase. In the collection described in the previous section, 69 different kinases were perturbed, associated with an increase or decrease in their activity (up-regulation: 100, down-regulation: 259 cases) (Supplementary Fig. 2b).

After scaling the activities per experiment, we followed the benchmark pipeline in the decoupler python package (See methods section “*Perturbation benchmark procedure*”) to compare the performance of all combinations of the computational methods and kinase-substrate libraries described above. Similar to previously proposed^31^, we also tested how the number of targets for each kinase might affect the performance, by taking the measured number of targets in an experiment as the kinases’ activity values. Additionally, we used a randomized kinase-substrate library as a baseline for performance. In this library, the phosphorylation sites reported in PhosphoSitePlus, as one of the most commonly used libraries for kinase activity inference, were shuffled and randomly assigned to an upstream kinase.

To evaluate the performance for each combination, all inferred kinase activities across experiments were sorted by their activity scores and the area under the receiver operating characteristic curve (AUROC) was calculated based on the perturbation information for each experiment (See methods section “*Perturbation benchmark procedure*”). Additionally, we calculated the scaled rank for each kinase within its experiment by dividing the rank of the perturbed kinases by the total number of kinases with an inferred activity (See methods section “*Mean rank benchmark*”).

The z-score as implemented by RoKAI, as well as the sum, in combination with PhosphoSitePlus and PTMsigDB were the best performing methods with a median AUROC of 0.80 (Fig. 3a; Supplementary Fig. 3a). When comparing the performance of PhosphoSitePlus to the other libraries across all methods, we only observed significant differences for NetworKIN, iKiP-DB and the shuffled library, which overall had lower median AUROCs (adjusted p-value < 8.5×10^−5^, mean Wilcoxon statistic across libraries equal to 278). Furthermore, all kinase substrate libraries performed better than our randomized control (adjusted p-value < 4.7×10^−9^, mean Wilcoxon statistic across libraries equal to 288.5)(Supplementary Table 1). When comparing the computational methods, all methods, except for KARP, UQ and the number of targets, had a median AUROC of at least 0.7 in combination with PhosphoSitePlus (Fig. 3a). However, we could not identify significant differences between methods across kinase-substrate libraries (adjusted p-values > 0.05, mean Wilcoxon statistic across methods equal to 26)(Supplementary Table 2). As also previously observed^31^, the number of targets performed comparable well to other methods, indicating a bias for well-annotated kinases as perturbation targets in the experiments. However, in contrast to other methods, this metric also performed well for the shuffled network, as here the number of targets did not change compared to PhosphoSitePlus.

In addition to the AUROC, we compared the median scaled rank of the perturbed kinases across combinations.The scaled rank describes the quantile in which the perturbed kinase’s activity falls, with a lower value being better. This metric measures how likely a perturbed kinase is to be located at the extremes of the distribution of inferred activities in each experiment, evaluating the ability of a method to assist in kinase prioritization in a real-world experiment. For the z-score, the sum, KSEA, the normalized mean, PTM-SEA and the univariate linear model in combination with PhosphoSitePlus, PTMsigDB and OmniPath, the median scaled rank of the perturbed kinases was equal to or lower than the 0.25 quantile (Fig. 3b; Supplementary Fig. 3b). The number of targets performed comparable well to other methods again. As already described for the AUROC analysis, this indicates a bias for well-annotated kinases as perturbation targets in the experiments.

For the top performing library across methods, namely PhosphoSitePlus, we also investigated the performance in identifying perturbed kinases based on their activities for each kinase separately. For each kinase, we calculated its mean rank for every experiment in which it was perturbed and visualized it in a kinome tree (Fig. 3c). AURKB, ATM, AURKA, MAP2K2 and ALK ranked among the top 5 kinases on average whenever the respective kinase was perturbed. However, CSNK2A1 had an average rank of 70 whenever perturbed. To investigate whether these differences might be linked to the number of targets of a kinase or a study bias of the kinases, we correlated the number of targets and citations for each kinase with its rank, reflecting their potential study bias. However, we did not observe a strong correlation for either the number of targets (pearson correlation equal to 0.12, p-value equal to 0.42) or the number of citation (pearson correlation equal to -0.17, p-value equal to 0.28) and the rank of the kinases (Supplementary Fig. 3c-d).

In conclusion, perturbation-based benchmarking analysis revealed distinct clusters for method-resource combinations, and PhosphoSitePlus and PTMsigDB in combination with the z-score and the sum performed particularly well. In general, manually curated databases, such as GPS gold, PTMsigDB, PhosphoSitePlus and OmniPath demonstrated similar performances when assessed by both AUROC and scaled rank, with PhosphoSitePlus having the best performance overall. Additionally, methods like the z-score, KSEA, the sum, the normalized mean, or an univariate linear model clustered together and had the highest performance using both AUROC and scaled rank. The performance of the activity estimation differed between individual kinases, which could not be linked to any study bias.

**Figure 3.**
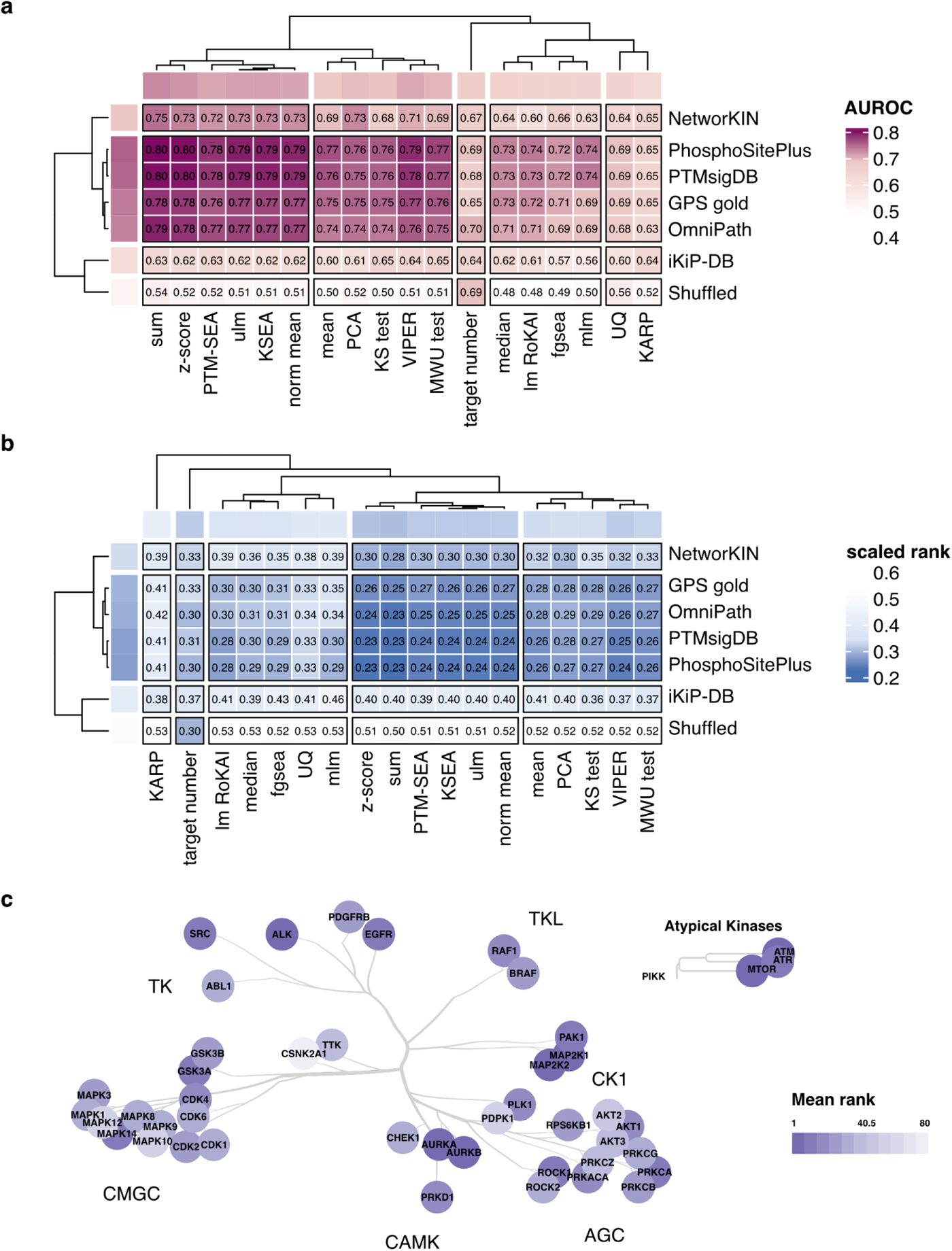
Systematic benchmark of kinase activity inference methods. **a** Predictive performance of methods for kinase activity inference in identifying perturbed kinases from phosphoproteomics data. Median AUROC for each kinase-substrate library - computational algorithm prediction. Hierarchical clustering was used on both libraries and methods. **b** Median scaled rank of the perturbed kinases’ inferred activity for each library-algorithm pair. The scaled rank is determined by taking the rank of the perturbed kinase in its experiment based on the activity, dividing it by the total number of kinases for which an activity was calculated. Hierarchical clustering was used on both libraries and methods. **c** Kinome tree of perturbed kinases each colored by its mean ranks across its perturbation experiments for the best performing kinase-substrate library PhosphoSitePlus.

### A complementary benchmarking approach using human tumor data to evaluate kinase activity inference

Even though perturbation experiments can be a helpful tool to benchmark kinase activity inference, they are typically constrained to a small set of well-studied kinases and may not adequately address off-target and indirect effects. Additionally, these experiments are usually performed in cell lines that lack the complexity of the tumor microenvironment observed in patients. Thus, we introduce a complementary benchmarking strategy that leverages multiple omics layers to construct a gold standard set of highly active or inactive kinases using human tumor profiling data from CPTAC.

CPTAC generated matched proteomics and phosphoproteomics data for the same set of tumors from ten cancer types^15^. Since overall phosphosite levels are commonly dependent on the levels of the corresponding host proteins, we first tested the correlation between phosphorylation sites and their host protein level in the CPTAC data. Phosphorylation sites showed similar distributions for correlation with their corresponding host proteins, with medians ranging from 0.36 to 0.47, for most cancer types (Supplementary Fig. 4a). However, colon (COAD), ovarian (OV), and pancreatic (PDAC) cancer, with median correlations of 0.32, 0.26, and 0.31, respectively, had distributions that were markedly lower. Since the lower correlation could suggest that these phosphoproteomics datasets may not be as robust, we excluded these three cancer types (two of these were also from an earlier round of CPTAC where the methodology was still being refined) from subsequent analyses. Therefore, this benchmarking approach focuses on breast cancer (BRCA)^53^, clear cell renal carcinoma (CCRCC)^54^, glioblastoma (GBM)^55^, head and neck squamous cell carcinoma (HNSCC)^56^, lung adenocarcinoma (LUAD)^57^, lung squamous cell carcinoma (LSCC)^58^, and uterine corpus endometrial carcinoma (UCEC)^59^ (Fig. 4a).

To use this resource to benchmark kinase activity inference in tumors, we started with the hypothesis that tumors with the highest kinase protein levels would have the highest kinase activities whereas tumors with the lowest levels would have the lowest activities. To test this hypothesis, we used the z-score as implemented by RoKAI in combination with kinase substrates from PhosphoSitePlus, as this was the best performing combination in the perturbation benchmark implemented above. Across all cancer types, z-scores were significantly higher in tumors in which the corresponding kinase protein levels were in the top 5% compared to those in the bottom 5% (Fig. 4b). These results support the use of the upper and lower tails of the abundance distribution for individual kinases to identify samples with high and low kinase activities, respectively. Accordingly, we defined a gold standard set of kinase-tumor pairs to benchmark different methods of kinase activity inference (Fig. 4c-d, methods section “*Development of a tumor-based benchmark*”). Briefly, we applied ROC analysis to determine how well each method distinguishes between kinase-tumor pairs in the gold standard positive set (top 5%) and those in the gold standard negative set (bottom 5%). To mitigate cancer type differences while maintaining the power gained from including all seven cancer types, the gold standard kinase-tumor pairs were selected for each kinase in each cancer type separately, and the inferred kinase activity scores were converted to z-scores within each cancer type. The ROC analysis was then performed on the combined set of kinases and cancer types. To also assess the stability of the results from the ROC analyses, we subsampled 80% of the gold standard set 1,000 times.

With AUROC values of ∼0.66-0.67, 10 out of 18 methods, including KSEA and PTM-SEA, performed similarly to the best performing method, namely the z-score (Fig. 4e). With AUROC values of ∼0.64-0.65, 7 of the 8 remaining methods were only marginally worse than the 10 best performing methods, whereas the performance for KARP was significantly lower than any of the other methods and barely better than the controls (Fig. 4e, Supplementary Fig. 4b). Similar trends were observed when different thresholds (top/bottom 2.5% and 10%) were used to select positive and negative pairs for the gold standard sets (Supplementary Fig. 4c-f) and when other prior knowledge substrate databases were tested (Supplementary Table 3) except for iKiP-DB, where the overall performance range was substantially lower (0.53-0.54). Thus, the performance results obtained from this new benchmarking approach are consistent with the established perturbation-based approach. For both, simple methods such as the mean or z-score perform similarly to more complex methods such as PTM-SEA.

**Figure 4.**
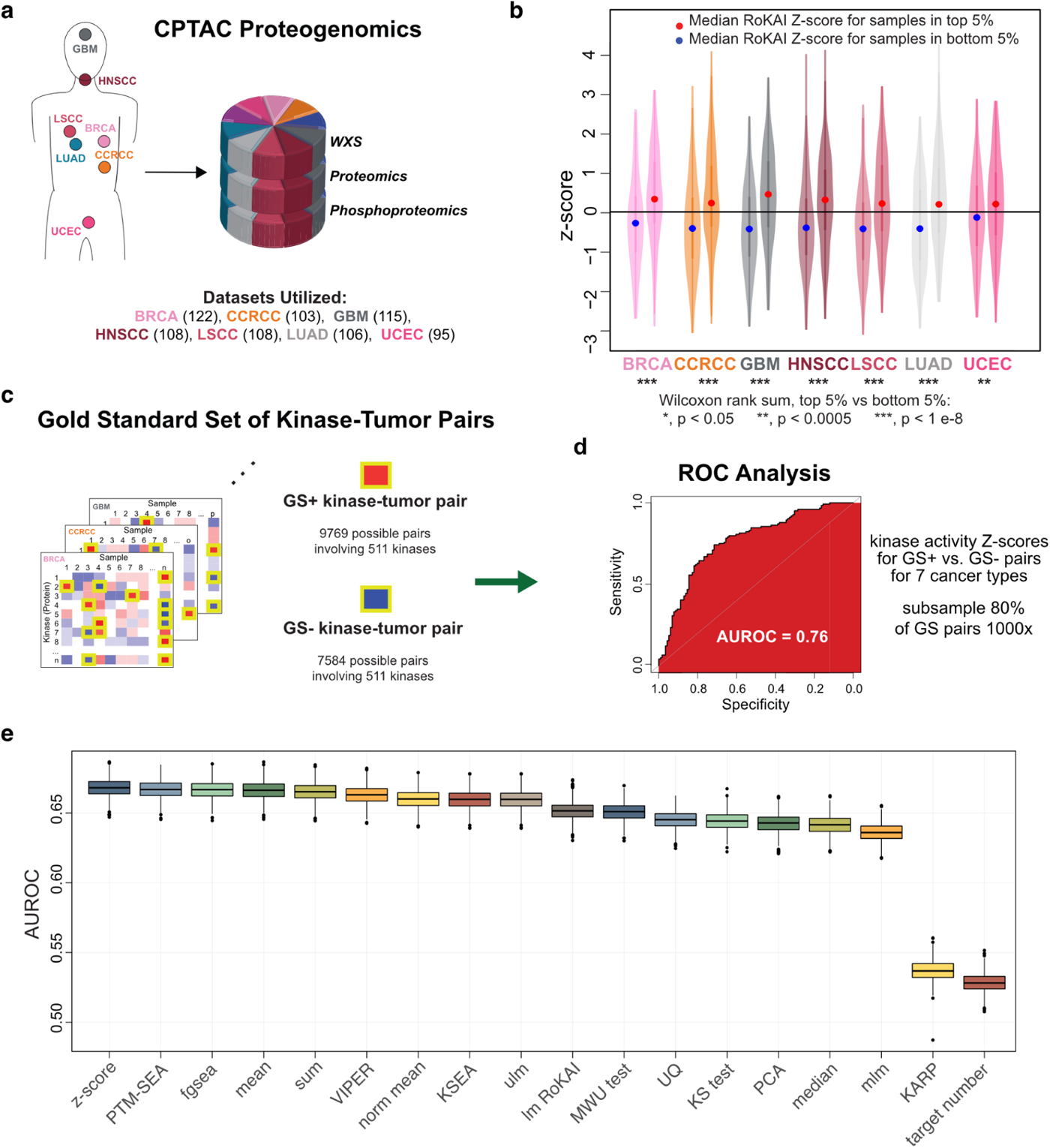
Development of a human tumor-based benchmarking approach using multi-omics data from CPTAC. **a** Summary of CPTAC datasets used in the current study. Cancer types included here are breast (BRCA), kidney (clear cell renal cell carcinoma: CCRCC), brain (glioblastoma: GBM), head and neck (HNSCC), lung (adenocarcinoma: LUAD and squamous cell carcinoma: LSCC), and uterine/endometrial (UCEC) cancers. The number of patients for each cancer type is shown in parentheses. **b** Inferred kinase activity scores from the phosphoproteomics data are significantly higher in tumors with the highest relative levels of the corresponding kinase (top 5% according to the normal distribution of kinase protein levels) than in tumors with the lowest levels (bottom 5%). **c** Toy diagram demonstrating selection process for kinase-tumor pairs for the Gold Standard (GS) set to be used for benchmarking kinase activity. For each cancer type, the protein data for each kinase (rows) was used to identify samples (columns) with the highest protein levels for the GS+ (tred boxes highlighted in yellow) and the lowest protein levels for the GS-(blue boxes highlighted in yellow) sets. **d** Benchmarking approach. The activity scores for each kinase are first converted to Z-scores across all samples within a cohort, and receiver-operator curve (ROC) analysis is used to evaluate how well theses Z-scores distinguish between kinase-tumor pairs in the GS+ and GS-sets for all kinases across all cancer types pooled together. To account for variability, ROC analyses are repeated 1000x after randomly subsampling kinase-tumor pairs from each GS subset for which activity scores are available. **e** Comparison of all kinase activity inference methods using the CPTAC-derived benchmark. Boxplots show the distributions of AUROC scores from benchmarking analysis applied to 1000x random samples of 80% of the GS set (median is indicated by line in center, whereas upper and lower boundaries of box show upper and lower quartiles, respectively, and circles indicate outliers). AUROC: area under the receiver-operator curve.

### Normalizing sites to host protein levels reduces kinase activity inference performance

Since phosphorylation sites are often well correlated with and reflective of host protein levels (Supplementary Fig. 4a), one potential concern is that phosphorylation site differences driven by changes in host protein levels may mask differences due to changes in the activity of upstream kinases. Therefore, we utilized our benchmarking approach to determine whether normalizing phosphorylation site levels to host protein levels prior to calculating kinase activity improves kinase activity inference. Specifically, we focused on two types of normalization strategies: linear regression to account for host protein levels and subtraction of host protein log intensities from site log intensities (subtract). For linear regression, we used three approaches: a single global linear model for all sites vs. corresponding host proteins (global), separate linear models for all sites on each protein separately (protein), and separate models for each individual site (site). Unnormalized data provided the best input for maximal performance for all kinase activity inference methods (Fig. 5a). Data that was normalized with the global linear regression approach resulted in modest reductions in performance. For z-scores, the mean AUROC was 0.662 for the unnormalized data and 0.645 for the global linear regression normalized data (similar trends were observed for other methods). However, all other normalization strategies resulted in AUROCs lower than 0.62, suggesting that the adjustments imposed by these normalization approaches may be overcorrections that remove meaningful information in addition to potential confounding effects from the host protein levels. In support of the possibility that the protein data already contains some relevant signal, we found that host proteins of common targets of the same kinases showed significantly higher correlation than host proteins for targets of different kinases in the CPTAC data (Supplementary Fig. 5a).

### Adding predicted targets to boost kinase activity inference

One way to potentially improve accuracy and boost the number of sites that are considered for the activity inference of a given kinase is to add predicted or *in vitro* identified substrates from databases such as NetworKIN or iKiP-DB, respectively. To test this possibility, we compared the performance of each kinase activity inference method in both benchmarks when using just curated targets from PhosphoSitePlus to the performance obtained when complementing the curated targets with targets from the NetworKIN database or iKiP-DB.

Overall, the addition of targets from NetworKIN and iKiP-DB boosted the number of kinases for which an activity could be inferred in both the CPTAC and the perturbation datasets and increased the number of kinases considered for evaluation by both benchmarks (Table 2). Furthermore, using the combined set of targets from PhosphoSitePlus and NetworKIN as substrates resulted in improved overall performance in the tumor-based benchmarking approach over using just known targets for all methods (Fig. 5b). Once again, using the z-score resulted in the best overall performance (AUROC = 0.68 for tumor-based), while the performance for KSEA (mean AUROC = 0.669) and PTM-SEA (mean AUROC = 0.679) was only modestly lower. However, this improvement could not be observed for the perturbation-based benchmarking approach (Fig. 5c). Additionally, adding iKiP-db targets led to a decrease of performance in both benchmarking approaches (Fig. 5b-c). Hereby, it is important to keep in mind that the evaluated kinases do not necessarily overlap between the two benchmarking approaches.

**Table 2.** Kinases covered in evaluation set.

| **Target set** | **Kinase total** | **Kinases per dataset** | **Kinases in evaluation set** | **Evaluated kinases per dataset** |
| --- | --- | --- | --- | --- |
| **Tumor benchmark:** |  |  |  |  |
| PhosphoSitePlus | 103 | 85 | 66 | 29 |
| PhosphoSitePlus & NetworkKIN | 157 | 134 | 104 | 51 |
| PhosphoSitePlus & iKiP-DB | 241 | 215 | 171 | 80 |
| **Perturbation benchmark:** |  |  |  |  |
| PhosphoSitePlus | 139 | 70 | 38 | Not applicable |
| PhosphoSitePlus & NetworkKIN | 196 | 108 | 50 | Not applicable |
| PhosphoSitePlus & iKiP-DB | 269 | 175 | 50 | Not applicable |

Since different kinases are likely contributing differently to the evaluation in our benchmarks, we also directly compared the performances for the different target sets after filtering to strictly the set of kinases evaluated for the PhosphoSitePlus target set itself in each cancer type (Fig. 5d-e). For this set of kinases, which are more likely to be well-studied kinases with better characterized substrates, the performance boost obtained from the combined target set was markedly higher for the tumor-based benchmark (mean AUROC 0.70 in the combination vs 0.67 for PhosphoSitePlus alone) and slightly higher in the perturbation-based benchmark (mean AUROC = 0.80 for the combination vs. 0.79 for PhosphoSitePlus), while the performance for the iKiP-DB set was still lower but not as poor as when considering all kinases (mean AUROC = 0.65 tumor-based; mean AUROC = 0.78 perturbation-based).

**Figure 5.**
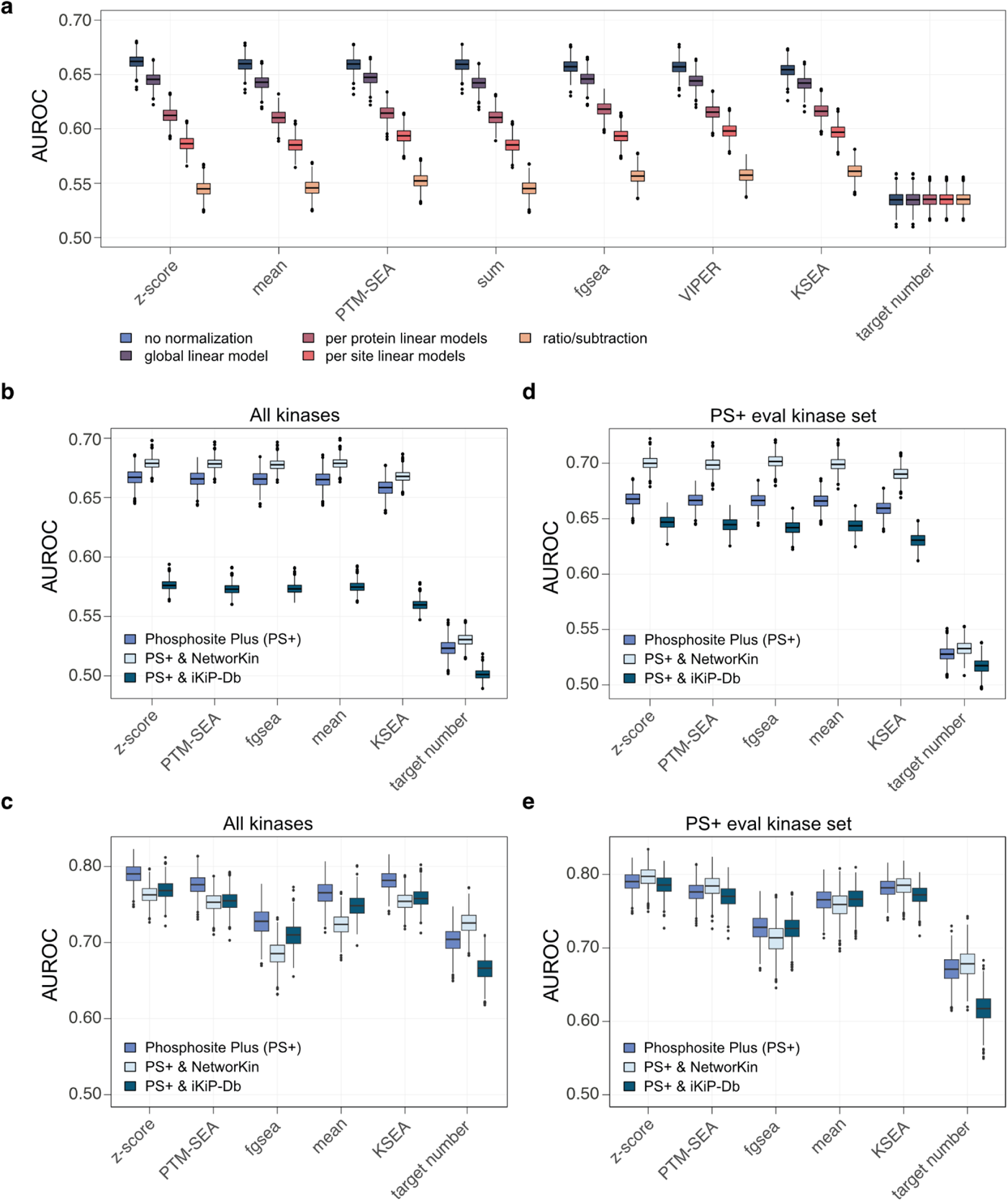
Utilizing benchmarking to optimize kinase activity inference. **a** Normalizing phosphosite levels to host protein levels prior to kinase activity inference reduces performance in the tumor-based benchmark. **b** Kinase activity inference performance (AUROC) using the tumor-based benchmark approach when considering the target set combining known targets from PhosphoSitePlus with predicted targets from NetworKIN or iKiP-DB. **c** Kinase activity inference performance (AUROC) using the perturbation-based benchmark approach when considering the target set combining known targets from PhosphoSitePlus with predicted targets from NetworKIN or iKiP-DB. **d** Evaluation of performance using the tumor-base benchmark approach after filtering for kinases that could be evaluated using the PhosphoSitePlus target set in each cancer type. **e** Evaluation of performance using the perturbation-base benchmark approach after filtering for kinases that could be evaluated using the PhosphoSitePlus target set in each experiment. PS+: PhosphoSitePlus; AUROC: area under the receiver-operator curve.

Additionally, we compared the performance of PhosphoSitePlus alone with the combination of PhosphoSitePlus and targets predicted from the kinase library published by Johnson et al. and Yaron-Barir et al.^60,61^ as well as predicted targets using the large language models Phosformer^62^. However, adding targets predicted by the kinase library or Phosformer led to a decrease in performance for both benchmarking approaches (Kinase library: mean AUROC = 0.58 tumor-based; mean AUROC = 0.71 perturbation-based, Phosformer: mean AUROC = 0.57 tumor-based; mean AUROC = 0.72 perturbation-based)(Supplementary Fig. 5 b-c).

Based on this analysis, we chose to use the z-score for a combined set of known targets from PhosphoSitePlus and predicted targets from NetworKIN to infer kinase activity for latter analyses.

### Activating sites on kinases are better associated with inferred kinase activity than kinase protein levels

To determine if activating sites provide an additional layer of regulation beyond kinase protein levels, we also evaluated the correlation of these sites with corresponding host kinase protein levels in each cancer type. Using PhosphositePlus annotations in combination with manual review of the literature, we identified 787 phosphorylation sites on 280 kinases that are associated with activation of their host kinases (Fig. 6a). While activating sites were reasonably well correlated with kinase protein levels (median correlation ranged from ∼0.2 to 0.32, depending on cancer type), the correlation was noticeably lower for these activating sites than it was for the same number of randomly drawn non-activating sites and their host proteins (Fig. 6b). Empirical p-values reflecting how frequently 1000 samples of random sites showed correlation as low as or lower than activating sites ranged from 0.001 (CCRCC) to 0.18 (LUAD). On average, activating sites only showed correlation that was higher than random for ∼7% of the sample sets. Thus, changes in activating sites are less likely to reflect changes in the protein, supporting the hypothesis that they may be more likely to contribute to regulation of kinase activity beyond the regulation imparted by kinase protein levels themselves.

To further investigate this possibility, we compared the correlation of inferred kinase activity scores with kinase activating sites to their correlation with kinase protein levels. For the same set of kinases, activating sites showed significantly greater correlation with kinase activity than did protein levels (Fig. 6c, paired Wilcoxon rank sum test, p = 0.00093). However, the difference between mean Pearson r values was only 0.048, and the analysis presented in previous sections was based on the assumption that kinase activity was already largely dependent on kinase protein levels, which prompted a more nuanced investigation. The plot in Figure 6d shows the difference between the correlation with activity scores for each kinase activating site and its corresponding host protein (y-axis) vs. the correlation between the site and host protein (x-axis). When sites and host proteins are well correlated, the difference in the correlation for the two to the activity score tends to be smaller, but activating sites tend to show better correlation than protein when the correlation between the site and host protein is poor. In a direct comparison of activating site and the host protein correlations with activity, activating sites showed significantly greater correlation with kinase activity than did kinase protein levels (Fig. 6e, paired Wilcoxon rank sum, p = 0.02; difference in mean Pearson r values was 0.078) when the Pearson correlation between the site and host protein was less than or equal to 0.2, whereas there was no difference when the correlation was greater than or equal to 0.4 (Fig. 6f, paired Wilcoxon rank sum, p = 0.67; difference in mean Pearson r values was 0.009).

In essence, while kinase activity is largely driven by protein levels, particularly when activating sites are well correlated with those proteins, activating sites are better associated with kinase activity when the correlation is poor. Because of this observation, we also repeated the benchmarking analysis after using activating sites instead of kinase protein levels to define the gold standard sets. While this reduces the number of kinase-tumor pairs that are included in the gold standard set (Supplementary Table 4), similar results were obtained when substituting the gold standard sets defined using kinase protein levels with activating site gold standard sets (Supplemental Fig. 6a-d). Specifically, most methods had comparable performance using these benchmarks, and the z-score as implemented by RoKAI was consistently among the top performing methods.

**Figure 6.**
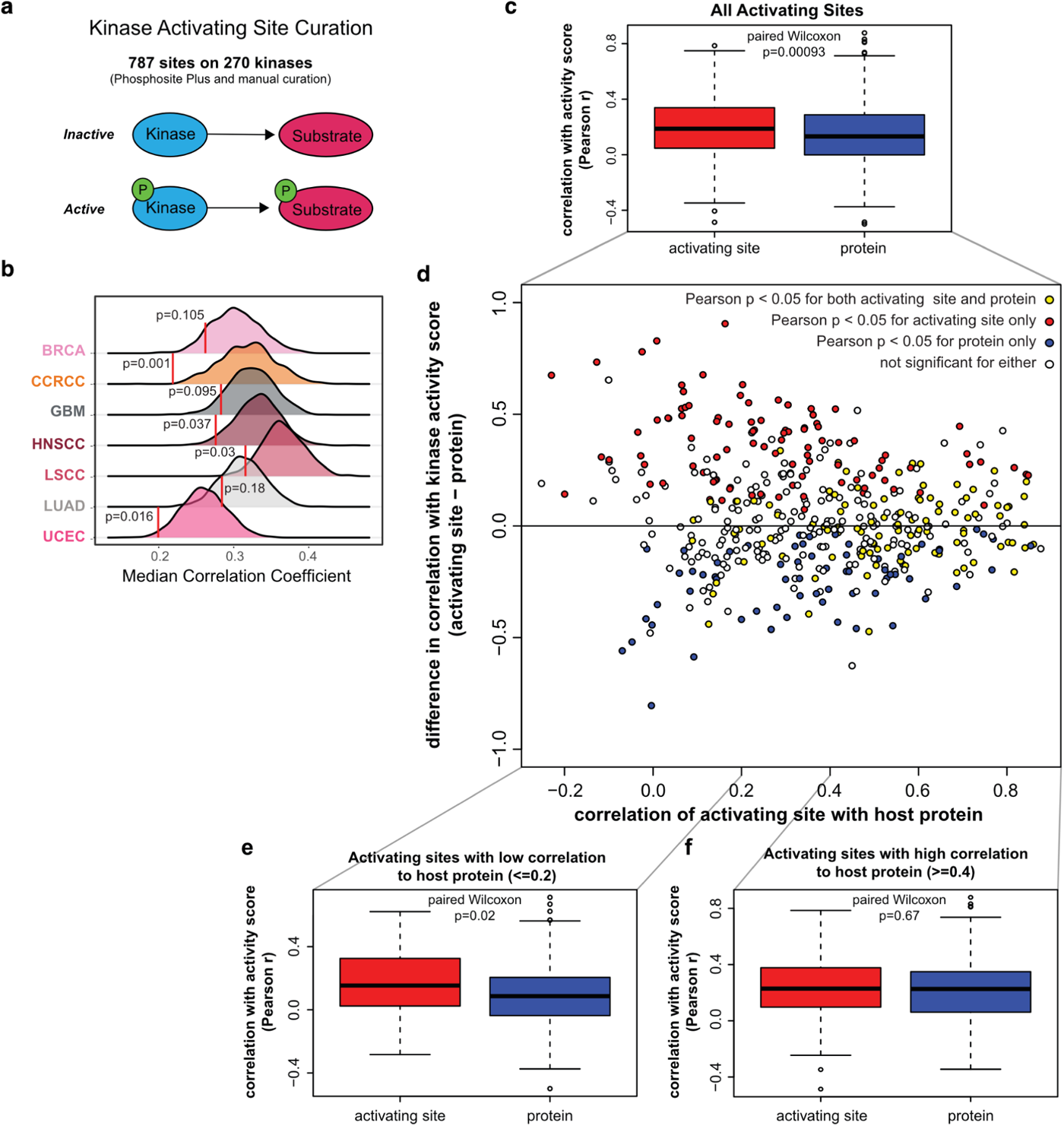
Activating sites on kinases are better associated with kinase activity than protein levels. **a** 787 potential activating sites on 270 kinases were identified from a combination of manual curation of the literature and of regulatory sites from PhosphoSitePlus. **b** Activating sites on kinases have lower correlation with their host proteins than sites selected at random. Red lines show the median Pearson correlation coefficient (r) for all activating sites with corresponding host proteins within each cancer cohort. Plots show the distribution of the median correlation coefficients for 1000 random samples of sites of the same size as the activating site set that are not activating sites, and the p-value indicates the fraction of random samples with median correlation coefficients equal to or less than the median correlation coefficients for the activating sites. **c** Activating sites on kinases show significantly higher correlation with activity scores than kinase protein levels do. Boxplots show the distributions of Pearson correlation coefficients between kinase activity scores and either activating site levels on the corresponding kinases (left) or kinase protein levels (right). (d-f) Activating sites with low correlation to host protein show better association with corresponding kinase activity scores than protein levels do, whereas protein levels and activating sites are similarly associated with activity when activating site-host protein correlation is high. **d** Scatter plot showing the difference between activating site and protein correlations with kinase activity scores (y-axis) vs. the Pearson correlation coefficient for the corresponding site with the corresponding host protein (x-axis). Each point represents an individual unique site in a single cancer type (all cancer types included in plot). **e** Boxplots show distributions for correlation coefficients for activity scores with activating site levels (left) or with protein levels (right) for kinases with low correlation (r <= 0.2) between the corresponding activating site and protein. **f** Boxplots show distributions for correlation coefficients for activity scores with activating site levels (left) or with protein levels (right) for kinases with high correlation (r >= 0.4) between the corresponding activating site and protein. For boxplots, the lines in the center show median values, whereas upper and lower boundaries of boxes show upper and lower quartiles, respectively, and circles indicate outliers).

### Kinase activity is a better marker for response to kinase inhibitors in cell lines than kinase protein levels

The primary objective of precision oncology is to identify tumors that would be good candidates for treatment with a specific drug. For kinase inhibitors, thus, we strive to target tumors with elevated kinase levels/activity. To determine if inferred kinase activity scores would serve as more effective indicators for response to treatment with a given kinase inhibitor than kinase protein levels, we utilized a systematic resource for testing drug response in the NCI60 collection of cell lines from the Genomics of Drug Sensitivity in Cancer (GDSC) project^63^. Since proteomics and phosphoproteomics data is available for many of the cell lines tested^64^, we correlated drug responses across these cell lines to protein levels of the corresponding kinase as well as to kinase activity scores. As an example, high CDK4 activity inferred using the z-score method from RoKAI and targets from PhosphoSitePlus combined with those from NetworKIN was associated with better response (lower AUC) to palbociclib, a CDK4/6 inhibitor, whereas higher protein levels tended to be associated with worse response (high AUC) (Fig. 7a). A systematic analysis of the association of drug response with target kinase measurements is shown in the dumbell plot in Figure 7b. Here, a lower AUC indicates better response; thus, the more negative the correlation, the better the association of a given kinase metric is with inhibitor response. The measurement best associated with response for the most inhibitor-kinase combinations tested (13/29) was the kinase activity score inferred using the combined set of known plus predicted targets, while kinase protein levels showed the greatest number of worst (highest) correlations (17/29). On the other hand, while the correlations with response for activity scores calculated using just known targets from PhosphoSitePlus was often similar to those for scores calculated using targets from both PhosphoSitePlus and NetworKIN, this metric was only the best associated with response for 4/29 drug-target combinations.

In conclusion, high kinase activity scores inferred using both known targets from PhosphoSitePlus and predicted sites from NetworKIN are better indicators of the likelihood for tumors to respond to treatment to the corresponding kinase inhibitor than kinase protein levels or activity scores inferred using known targets alone.

**Figure 7.**
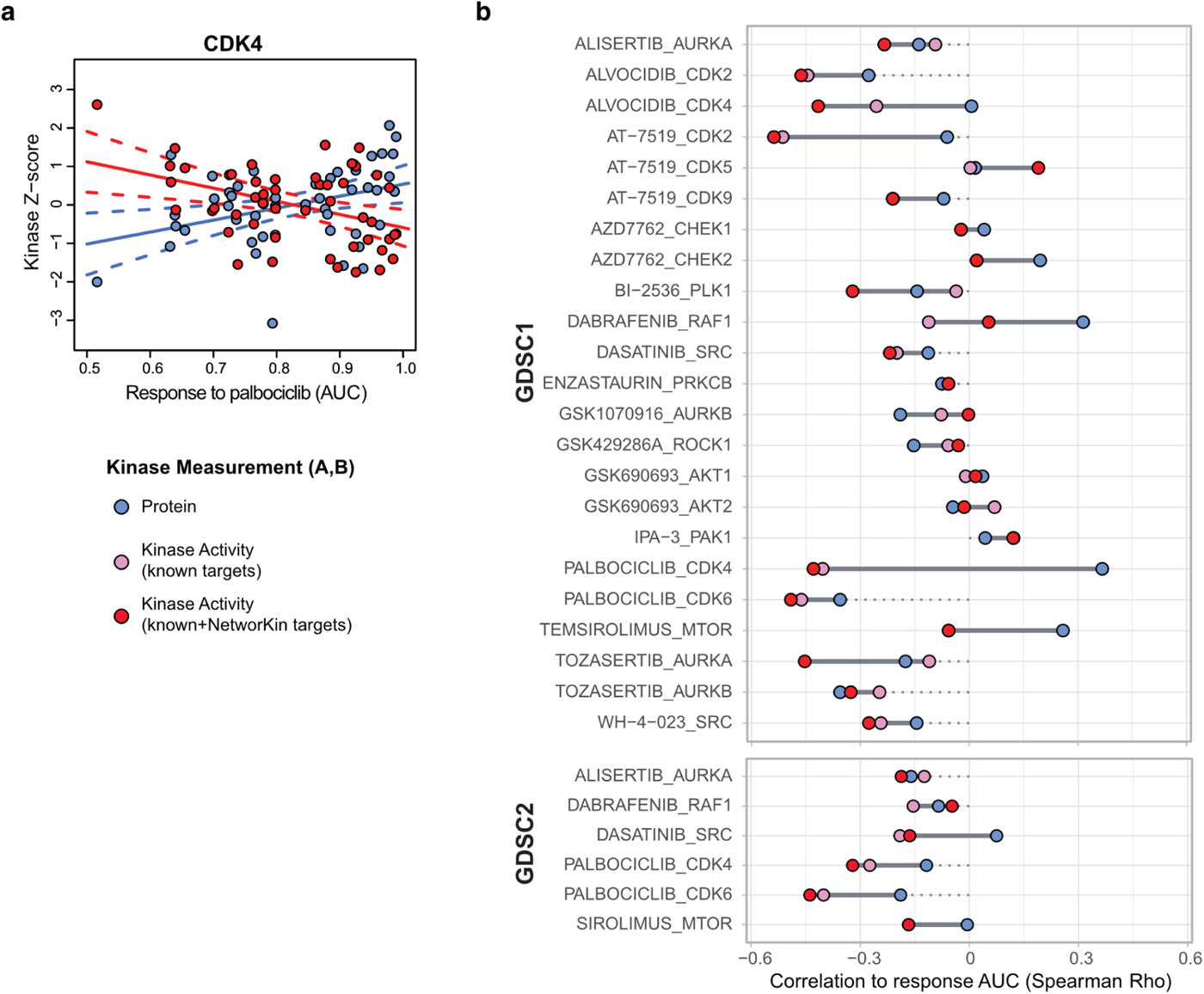
Inferred kinase activity scores provide a better indication for response to kinase inhibitor treatment than protein levels. **a** CDK4 activity scores are higher, whereas CDK4 protein levels are lower, in cell lines with better response to palbociclib, a CDK4/6 inhibitor. Scatter plot shows z-scores for CKD4 activity scores (red points) or protein levels (blue points) vs. palbociclib response data from the Genomics of Drug Sensitivity in Cancer project (GDSC1; lower AUC (area under curve) indicates better response). Each point represents a different NCI60 cell line. Solid lines are from fitted linear models associating activity (red) or protein (blue) with response AUC, and dotted lines show the 95% confidence interval for these models. **b** Dumbbell plot comparing Spearman Rho values for correlation of response to kinase inhibitors in NCI60 cell lines from the GDSC with levels of the corresponding kinase target protein (blue points) to correlations with the corresponding kinase activity score (red points show correlations with scores calculated using PhosphoSitePlus + NetworKIN targets and pink points show those for scores calculated using PhosphoSitePlus targets alone) levels. Correlations with drug response data from GDSC1 for each drug-kinase target pair are shown in the upper plot whereas correlations for GDSC2 data are shown in the lower plot.

## Discussion

The identification of deregulated kinases is a crucial component of biomedical research as they play central roles in signaling and serve as important drug targets. Therefore, computational tools have been developed to infer kinase activities from phosphoproteomics data. To interpret these findings accurately, it is important to critically evaluate the reliability and coverage of kinase-substrate libraries and prediction tools, as well as the impact of different computational algorithms.

In this paper, we present a systematic comparison of methods and prior knowledge resources for kinase target sets that can be used for kinase activity inference and introduce a new benchmarking strategy to identify the most reliable methods. Evaluation of all combinations of methods and prior target sets using the classical perturbation-based benchmarking approach identified that simpler computational methods like the mean or z-scores used by RoKAI or KSEA are comparable if not superior to more sophisticated methods like fgsea, or multivariate linear models. Additionally, manually curated target resources, especially PhosphoSitePlus, perform the best. Furthermore, we introduce a complementary benchmarking strategy based on multi-omics data from human tumors where again, simpler computational methods performed among the best. We next used both benchmarking approaches to test whether boosting PhosphoSitePlus targets with predicted targets from various resources could enhance kinase activity inference. Adding targets predicted by NetworKIN but not other prediction tools did improve performance noticeably in the tumor-based benchmark, but only modestly in the perturbation-based benchmark. Finally, we showcase the potential for kinase activity inference to inform precision oncology by identifying active kinases in a given sample.

We employed two complementary approaches for the evaluation of kinase activity inference methods because both have distinct strengths and limitations. The perturbation-based approach is straightforward and focuses on assessing the direct effect of the perturbation on the kinase’s activity. However, it is limited to usually well-studied kinases that have been experimentally perturbed and profiled by phosphoproteomics. Here, we also observed some bias for well-studied kinases, with the number of targets as a metric for kinase activity inference performing considerably well in the benchmark. Additionally, it could be confounded by downstream kinases or feedback regulations and usually does not account for off-target effects of drug perturbations. There have already been some efforts in identifying the target spectrum of kinase drugs^65,66^; however, these are usually linked to binding assays and as such do not necessarily reflect a change in activity. Ideally, further perturbation studies with very specific drugs would be useful to minimize the limitations of this benchmark approach. Specificity is critical because a major assumption of the perturbation-based approach is that non-targeted kinases are less affected by the perturbation and treated as negatives, which could result in decreased performance if they are in fact also affected by the perturbation. Experimental perturbations require experimental systems that lack the complexity of the tumor microenvironment present in humans. In contrast, the tumor-based benchmark approach aims to better account for this complexity, but it is primarily based on the assumption that tumors with high kinase protein levels have high kinase activity whereas tumors with low kinase protein levels have low activity. This may not necessarily be the case as kinases are also subject to post-translational regulation and abundance thus may not necessarily translate into activity^67^. Furthermore, while this approach does bypass some of the bias towards considering mainly well-studied kinases, allowing for the inclusion of under-studied kinases in the evaluation, kinase activity inference still requires having a set of reliable target sites for a given kinase in order to obtain scores to evaluate in the first place. Finally, while off-target effects of a perturbation are not an issue with this approach, it is still likely that, in a tumor with high kinase levels, downstream kinases will also be active. Because of the advantages and disadvantages of these complementary approaches, we recommend using both and determining the consensus between the two.

Aside from evaluating kinase activity inference methods and kinase target sets themselves, the benchmarking approaches described here can be used to evaluate other potential strategies for improving inference. For example, when proteomics data is available for the same samples that have been profiled with phosphoproteomics, as is the case for the CPTAC cohorts, a logical hypothesis would be that normalizing phosphosites to their host protein levels may reduce the possibility that substrate protein levels, which are often well correlated with phosphosite levels, confound kinase activity inference. However, all of the strategies we tested for normalizing sites to host protein levels resulted in decreased performance in the tumor-based benchmark, suggesting that there may be signal present that is associated with kinase activity in the protein data already and that removing this signal impairs our ability to accurately infer that activity. In support of this hypothesis, we observed that the host proteins of common targets of the same kinase already show higher correlation than the targets of kinases from different groups, possibly because many kinases and their substrates may lie in common pathways. Alternatively, the noise present in both the phosphosite and protein data may be amplified by the normalization process; thus, normalization may actually be decreasing the signal-to-noise ratio. Both of these possibilities are supported by our observation that the reduction in performance is most severe when attempting to make this adjustment on a site by site basis (subtracting or using the per site linear model) but is minimal when the adjustment is made by considering the data as a whole (global linear model). The development of normalization strategies that mitigate these issues may allow us to better account for the effects of substrate protein levels when inferring kinase activity in the future.

By assessing the impact of utilizing predicted targets for kinase activity inference on performance, benchmarking may also allow us to evaluate the biological relevance of those predictions. Despite good performance for PhosphoSitePlus or other curated libraries, current kinase-substrate libraries are limited in their coverage of both kinases and phosphorylation sites. As a result, much of the information measured in phosphoproteomics cannot be used for the prediction of kinases activities and activity inference is restricted to a certain set of kinases, limiting the interpretation of the results. To bridge these gaps, a variety of substrate prediction tools have been introduced. However, besides NetworKIN, the tools tested here did not improve performance. This might be due to the fact that these tools solely focus on the amino acid sequence, neglecting context-dependency and the regulatory environment of kinases. As such, incorporating factors such as protein-protein interactions (PPI), subcellular localization, and the presence of inhibitors or activators could be crucial components to identify direct targets of kinases and ultimately improve predictions for kinase activities. Despite being the oldest of the prediction methods we tested, NetworKIN likely provided the best performance because the predictions are informed by PPI in addition to sequence homology to known targets. Unfortunately, NetworKIN is not actively maintained, and the substrate predictions used for our analysis were obtained by mapping the sites in our data to predicted sites downloaded from the website; predicted targets would have been overlooked if they could not be mapped properly. More modern kinase substrate prediction methods that use current protein sequence databases (or that are database agnostic) and incorporate biological context are likely to further improve performance, and our benchmarking tools should provide the means to evaluate how well they do so.

Kinase activity inference also has the potential to inform precision oncology. Tumors with high kinase activity scores may be candidates for therapy with the corresponding kinase inhibitors. Our cell line analyses indicated that the method and target site set that were among the best performing in the benchmarks also generated scores that were consistently best associated with inhibitor response. However, this analysis was performed using data from cell lines, and further studies will need to be carried out to confirm these trends in more complex biological systems. Furthermore, performing phosphoproteomic profiling of tumors to identify kinases to target is not practical in a clinical setting. A simpler and more quantitative assay using specific peptides that are representative of kinase activity would be more ideal. Our comparison of activating sites on kinases to kinase protein levels suggests that the best approach to select markers for kinase activity may be to consider each kinase on an individual basis; for some kinases, the protein levels may provide a sufficient readout for activity whereas activating sites or specific target sites that are well correlated with kinase activity may be better candidates for other kinases.

In summary, we performed a comprehensive comparison and identified PhosphoSitePlus in combination with targets from NetworKIN and z-score implemented by RoKAI as the best performing approach for kinase activity inference. Furthermore, both benchmarking approaches are available in the R package benchmarKIN (https://github.com/saezlab/benchmarKIN). The package includes all necessary data for both approaches and provides vignettes demonstrating how to use the benchmarking approaches for evaluating kinase activity inference. This will help to simplify the process of evaluating novel methods or other kinase-substrate resources.

## Methods

### Prior knowledge resources

We collected kinase-substrate interactions from multiple prior knowledge resources, including PhosphoSitePlus^2^, PTMsigDB^19^, the gold standard set used to train GPS 5.0 (GPS gold)^37^, OmniPath^38^, iKiP-DB^27^ and NetworKIN^29^. PhosphoSitePlus is a well established kinase-substrate resource containing curated, experimentally observed kinase-substrate interactions. We downloaded the kinase-substrate interactions from the PhosphoSitePlus website (https://www.phosphosite.org, accessed: 19/04/2023), which were filtered to retain only those reported in humans. Additionally, we sourced PTMsigDB signature sets, a collection of site-specific signatures for perturbations, pathways and kinases curated from literature. It is based on PhosphoSitePlus and considers information from NetPath, WikiPathways and LINCS. In newer versions it also includes signatures from iKiP-DB (in vitro Kinase-to-Phosphosite database). The v2.0.0 kinase sets tailored for humans were downloaded from the proteomics broadapp website (https://proteomics.broadapps.org/ptmsigdb/, accessed: 20/12/2023), and all kinase signatures were extracted, except for the ones from iKiP-DB as this was tested as a separate resource. GPS gold was downloaded from the supplementary files of the original publication (https://www.ncbi.nlm.nih.gov/pmc/articles/PMC7393560/#s0065). We next extracted all kinase-substrate interactions stored in OmniPath using the get_signed_ptms function from the *OmniPathR v3.11.1* package and filtered the interactions for phosphorylation events. OmniPath is a meta-resource, integrating information from over 100 different resources about intra- and inter-cellular signaling, including information about kinase-substrate interactions coming from PhosphoSitePlus, MIMP^43^, dbPTM^40^, HPRD^41^, SIGNOR^23^, PhosphoNetworks^45^, phospho.ELM^24^ and Li2012^42^, as well as ProtMapper^68^ and KEA3^34^. However, here all interactions exclusively reported by ProtMapper or any of the KEA3 libraries were excluded. NetworKIN was downloaded from the networKIN website (http://netphorest.science/download/networkin_human_predictions_3.1.tsv.xz) and filtered for interactions with a NetworKIN score equal to or higher than five. The NetworKIN database contains precomputed kinase-substrate interactions for the human phosphoproteome reported in the KinomeXplorer-DB. Kinase-substrate interactions were computed using the NetworKIN algorithm which integrates consensus substrate motifs with context modeling. Lastly, we included the iKiP-DB which contains kinase-substrate interactions identified based on a large-scale in vitro kinase study for over 300 human protein kinases. Kinase-substrate interactions were sourced from the supporting information of the corresponding publication (https://pubs.acs.org/doi/suppl/10.1021/acs.jproteome.2c00198/suppl_file/pr2c00198_si_007.zip). All kinase-substrate resources were processed to a common format, with kinases and target proteins expressed as human gene names, and filtered for kinases reported by kinhub or gene ontology (GO) terms (GO:0016301). Moreover, we extracted information such as the phosphorylated amino acid, its position within the protein, and, where available, the 15-mer or 11-mer (in case of NetworKIN) sequence surrounding the phosphorylated amino acid from all resources. For mapping the phosphorylation sites from the experiments to the phosphorylation sites in the kinase-substrate libraries, we used the combination of HGNC symbols with the flanking sequence of the phosphorylation site (15mer or 11mer), whenever applicable, or the combination of HGNC symbol with amino acid type and location information.

### Computational methods for kinase activity inference

For kinase activity inference, we select multiple published methods including fgsea^48^ (fast gene set enrichment analysis), KARP^49^ (Kinase activity ranking using phosphoproteomics data), KSEA^21,22^ (kinase set enrichment analysis), the linear model and z-score as implemented in RoKAI^31^ (Robust kinase activity inference), PTM-SEA^19^ (PTM-signature enrichment analysis) and VIPER^51^ (Virtual Inference of Protein-activity by Enriched Regulon analysis). Additionally, we included the following methods: Kolomogorov-Smirnov test, Mann-Whitney-U test, the mean, the median, the multivariate linear model, normalized mean and univariate linear model as implemented in decoupler^50^, principal component analysis, the sum and the upper quantile. A small description of all computational methods can be found below. For more detailed information please refer to the original publications.

***fgsea.*** Fast gene set enrichment infers kinase activities using a weighted running sum approach. It first ranks molecular features per sample and calculates an enrichment score by walking down the list of features, increasing a running sum statistic when a feature is part of the target set and decreasing when it is not. The magnitude of the increment depends on the correlation of that feature with the phenotype. The enrichment score is then the maximum derivation from zero. Here we used the implementation of fgsea from the *decoupler package v2.8.0*.

***KARP.*** KARP calculates a K-score which consists of the ratio of the sum of molecular features of the targets of a kinase and the sum of molecular features of all phosphorylation sites. This is then corrected for the imbalance in known targets by multiplying the square root of measured targets of a kinase divided by the total number of known targets of that kinase in a given resource. The K-score was implemented as described in Wilkes et al..

***KSEA.*** Kinase set enrichment analysis calculates a z-score to assess the difference between the mean of the molecular features of known targets of a kinase and the mean of molecular features of all phosphorylation sites, adjusted by the square root of the number of identified targets and the standard deviation of the molecular features of all phosphorylation sites. We implemented the KSEA method as described in the KSEAapp.

***Kologomorov-Smirnov.*** The Kologomorov Smirnov test compares the running sum of targets and non-targets of a kinase. Molecular features are again ranked and the running sum statistic increases if the feature is part of the target list. The increments hereby are always the same (in contrast to fgsea). To run the Kologomorov-Smirnov test we used the *ks.test* function from the *stats package v4.3.3* and used the negative logarithm of the p-value as the final score.

***Linear model - RoKAI.*** The linear model described by RoKAI simultaneously models the molecular readouts of all molecular features for all kinases. Thereby the phosphorylation site is modeled as the sum of activities for all linked kinases and the weights of non-targets are thereby set to zero. The kinase activity is inferred using least squares optimization including ridge regularization. To run the linear model we transcoded the original implementation from MATLAB to R.

***Mann-Whitney-U.*** The Mann-Whitney U test, also known as the Wilcoxon rank-sum test, compares the ranks of the molecular features between targets and non-targets of a kinase. All phosphorylation sites are ranked together based on their molecular features and the U-statistic is calculated based on the sum of ranks for targets and non-targets. To run the Mann-Whitney-U test we used the *wilcox.test* function from the *stats package v4.3.3* and used the negative logarithm of the p-value as the final score.

***mean.*** The mean refers to the average level of the molecular features of all target sites of a kinase.

***median.*** The median refers to the middle value of the molecular features of all target sites of a kinase when these features are ranked in order.

***multivariate linear model.*** The multivariate linear model as implemented in decoupler simultaneously models the molecular readouts of all molecular features for all kinases. Thereby the phosphorylation site is modeled as the sum of activities for all linked kinases and the weights of non-targets are thereby set to zero. Here, we used the *run_mlm* function from the *decoupler package v2.8.0*.

***normalized mean.*** For the normalized mean, random permutations of target features are performed and a random null distribution of means is obtained. For the normalized mean the average level of the molecular features of all target sites of a kinase is then subtracted by the mean of the random null distribution and divided by the standard deviation of the random null distribution. Here we used the implementation of the normalized mean from the *decoupler package v2.8.0*.

***principal component analysis.*** Principal component analysis is performed across samples using only the molecular features of the targets of a certain kinase. The score of that kinase then represents the variance explained by the first principal component. Here we used the *prcomp* function from the *stats package v4.3.3* without scaling.

***PTM-SEA.*** PTM-SEA calculates an enrichment score as described for fgsea. Additionally, it calculates a normalized enrichment score using random permutations of target features. To run PTM-SEA we used the *run_ssGSEA2* function from the *ssGSEA2 package v1.0.1*. We used the default settings and increased the maximum number of targets to 100,000.

***sum.*** The sum refers to the summed up levels of the molecular features of all target sites of a kinase. Here we used the implementation of the sum from the *decoupler package v2.8.0* which is also able to take interaction weights into account. These were all set to one here.

***univariate linear model.*** The univariate linear model as implemented in decoupler models the molecular readouts of all molecular features per kinase. The weights of non-targets are thereby set to zero and the obtained t-value from the fitted model represents the activity of the kinase. We used the *run_ulm* function from the *decoupler package v2.8.0*.

***upper quantile.*** The upper quantile represents the value below which 75% of the molecular features of all target sites of a kinase fall.

***VIPER.*** VIPER estimates kinase activities through a three-tailed enrichment score calculation based on the ranking of all phosphorylation sites and the targets of a kinase based on their molecular features. Finally, a normalized enrichment score is estimated using random permutations. For the implementation of VIPER we used the *decoupler package v2.8.0*.

***z-score - RoKAI.*** The Z-score as implemented in RoKAI calculates the mean of the molecular features of the known targets of a kinase and adjusts it by the square root of the number of identified targets for the kinase and the standard deviation of the molecular features of all phosphorylation sites. To run the z-score we transcoded the original implementation from MATLAB to R.

### Processing of kinase perturbation dataset

***Hernandez-Armenta.*** To evaluate the performance of different kinase inference methods combined with different prior knowledge networks, we obtained a curated set of 93 knockdown and overexpression perturbation experiments previously used by Hernandez-Armenta et al. We downloaded the processed datasets from Zenodo (https://zenodo.org/records/5645208). These datasets contain quantile-normalized log fold-changes from publicly available studies covering 27 different kinases regulated in 103 different perturbation settings.

***Hijazi.*** Additionally, we collected the perturbation datasets from Hijazi et al. containing phosphosite log fold-changes from 61 kinase inhibitors perturbation experiments in two different cell lines (HL60, MCF7). To identify the detected phosphosites we used the primary UniProt Accession number reported in the original data for each phosphopeptide. We filtered out phosphosites whose identification false discovery rate was greater than 0.05, as this was the threshold used in the original publication. Finally, for each perturbation experiment we generated a ranked list of peptides with their reported log fold-change. Peptides with multiple phosphorylation sites were split and in case of ambiguity we selected the fold change with a higher significance for the phosphorylation sites. We then collected the targets for each kinase inhibitor based on the Therapeutic Target Database (https://idrblab.net/ttd/), if available. The full list of targets and references can be found in Supplementary Table 5.

### Kinase activity inference

Kinases activities were estimated based on the log fold-change of the direct target phosphorylation sites for each kinase. To select the direct targets of a kinase we used different kinase-substrate libraries in combination with multiple computational methods, which are described in section *“Prior knowledge resources”* and section *“Computational methods for kinase activity inference”* in more detail. Using each combination, we inferred kinase activities for each experiment, considering only kinases with at least five measured target phosphorylation sites.

### Comparison of activity scores

We performed pairwise Pearson correlation to compare the inferred kinase activities scores from each prior-method combination for each experiment. We then performed hierarchical clustering on the mean Pearson correlation across experiments between all method-prior combinations.

### Perturbation benchmark procedure

Kinase activity scores obtained as described above were first multiplied by the sign of the perturbation (knockout: negative, overexpression: positive) for each perturbation experiment. To account for variability across experiments, the scores were scaled by dividing each score with the absolute maximum score of the experiment. Each method-resource combination was then evaluated using the get_performance function from the *decoupler v1.6.1* Python package.

Therefore the activity scores matrix (rows: experiments, columns: TF activities) is flattened across experiments into a single vector and missing values for activity scores are removed. Due to differences in class imbalance across networks, a downsampling strategy is employed within the benchmark. For each permutation, an equal number of positive and negative classes are randomly selected to calculate the area under the Receiver Operating Characteristic (AUROC) and Precision-Recall Curve (AUPRC) metrics. This process is repeated 1,000 times per network, obtaining distributions of performance measurements. To compare the performance across kinase-substrate libraries and across methods, we aggregated the median AUROC for each library across methods and vice versa. We then compared the median AUROCs between libraries and methods using a Wilcoxon test and performing p-value adjustment using Benjamini-Hochberg.

### Mean rank benchmark

We calculated the mean rank across experiments for each method-resource combination. Kinase activity scores of each experiment were multiplied by the sign of the perturbation (knockout: negative, overexpression: positive) and ranked by their activity. The ranks of the perturbed kinases were then selected and scaled by dividing the rank with the total number of inferred kinase activities. This scaled rank describes the quantile in which the perturbed kinase was found based on its activity, with a lower value being better. We then calculated the mean scaled rank of perturbed kinases across all experiments. Additionally, we calculated the rank for each perturbed kinase across its perturbation experiments. We then used CORAL (http://phanstiel-lab.med.unc.edu/CORAL/) to visualize the rank of each kinase across all method combinations from PhosphoSitePlus. Within the kinome tree, only perturbed kinases which were captured at least once are shown.

### Evaluation of study bias

The mean rank for each perturbed kinase was compared to their study quantity. Therefore, a table containing the PubMed IDs linked to each gene (gene2pubmed.gz) was downloaded from the National Institute of Health (https://ftp.ncbi.nih.gov/gene/DATA/). We calculated the number of unique PubMed IDs for each perturbed kinase and performed Pearson correlation to assess the relationship between the rank and the number of PubMed references of each kinase using the cor.test function from the *stats package v4.3.3* in R. Additionally, we assessed the relationship between the rank and the number of reported targets in PhosphoSitePlus for each kinase using the cor.test function from the *stats package v4.3.3* in R.

### Addition of predicted kinase-substrate relationships

To evaluate the addition of predicted kinase-substrate relationships we combined the kinase-substrate interactions reported by PhosphoSitePlus with predicted targets from the Kinase Library and Phosformer. The predicted interactions were selected as follows.

***Kinase Library.*** We calculated the percentile score for each substrate in the datasets based on the position-specific score matrices (PSSMs) derived from the positional scanning peptide array for all Serine/Threonine and Tyrosine kinases as presented by Johnson et al. and Yaron-Barir et al.^60,61^. Following Johnson et al., we then assigned the highest scoring 15 kinases based on their percentile scores as upstream regulators for each phosphorylation site.

***Phosformer.*** To obtain the protein language model predictions, we applied the Phosformer model^62^ (https://github.com/esbgkannan/phosformer to every phosphosite in our datasets to obtain the probability of upstream regulation for every kinase in the reference kinases list (https://github.com/esbgkannan/phosformer/blob/main/data/reference_human_kinases.csv). Following the threshold applied for the kinase library, we assigned the highest scoring 15 kinases as upstream regulators for each phosphorylation site.

### Development of a tumor-based benchmark

The data used to establish the tumor-based benchmark is the version of the CPTAC data harmonized across ten cancer types using the BCM pipeline described in Li et al.^15^. Based on the analysis of site-host protein correlations presented in Figure 4b, we chose to focus on data from the breast cancer (BRCA)^53^, glioblastoma (GBM)^55^, clear cell renal carcinoma (CCRCC)^54^, head and neck squamous cell carcinoma (HNSCC)^56^, lung squamous cell carcinoma (LSCC)^58^, lung adenocarcinoma (LUAD)^57^, and uterine corpus endometrial carcinoma (UCEC)^59^ studies. For each cancer type, the protein data for each kinase was used to identify samples for the Gold Standard positive (GS+) (those in the top 5% relative to the normal distribution of the protein levels; z-score > 1.645) and negative (GS-) (bottom 5% relative to the normal distribution; z-score < 1.645) sets after filtering out proteins with fewer than 30 measurements and with variance < 0.1. Alternative GS sets were also established using the top 2.5% (|z| > 1.96), 10% (|z| > 1.282), and 15% (|z| > 1.036; not analyzed here, but presented as an option in the benchmarKIN package). To establish alternative GS sets using activating sites on kinases, the same thresholds were applied to kinases phosphosite levels for the sites defined in the Analysis of activating sites on kinases methods section instead of kinase protein levels.

For benchmarking using these GS sets, we first median centered the log2 MS1 intensity data for each site within each cancer type to use as input for calculating kinase activity scores as described above (*Computational methods for kinase activity inference*). To ensure that there was no leakage from the data used to define the GS sets to the data used to calculate kinase activity scores, phosphorylation sites for the respective kinase were removed from each kinase target set from each prior knowledge kinase-substrate resource prior to kinase activity inference. The activity scores for each kinase were first converted to z-scores across all samples within a cohort, and receiver-operator curve (ROC) analysis was used to evaluate how well the z-scores distinguished between kinase-tumor pairs in the GS+ and GS-sets for all kinases across all cancer types pooled together. To account for variability, ROC analyses were repeated 1000x after randomly subsampling 80% of the kinase-tumor pairs from each GS subset for which activity scores are available. Functions (R code) for using this benchmarking approach with any of the GS sets described here given kinase activity scores calculated from the same CPTAC datasets are available in the benchmarKIN R package.

### Normalization of phosphosite levels to protein levels

To normalize the phosphosite data to host protein levels, we first filtered the protein and phosphosite log2 MS1 intensity data to remove proteins and sites with fewer than 30 measurements. Sites lacking measurements for respective host proteins were then removed. We then employed multiple strategies to normalize the level of each site in each dataset to the corresponding values of the host protein. Thus, the unnormalized data used for this analysis is different from the data used for the corresponding analyses presented in Figure 4. The first normalization strategy involved subtracting the log2 MS1 intensity of the protein in a given sample from the log10 MS1 intensity of the site. The remaining strategies relied on using the residuals from linear regression of the sites to their host proteins. We used three different types of models for the linear regression normalization: a single global linear model for all sites vs. corresponding host proteins (global), separate linear models for all sites on each protein separately (protein), and separate models for each individual site (site). Site median-centered data was used as input for kinase activity inference using PhosphoSitePlus targets as described above (*Computational methods for kinase activity inference*) and evaluation was carried out using the tumor-based benchmark.

### Analysis of activating sites on kinases

A list of manually curated activating sites from the literature was updated with sites annotated as promoting kinase activity in the regulatory sites file downloaded from PhosphoSitePlus^2^ in March 2022. For all activating sites with host proteins measured in the CPTAC data, Pearson correlation coefficients between the sites and corresponding host proteins were calculated for each cancer type. To determine if these correlations were lower than expected by chance, empirical p-values were calculated by randomly sampling an equal number of sites in each dataset and calculating the Pearson correlation between sites in these samples and the corresponding host proteins. The p-value is the fraction of these samples that had median correlation coefficients that were equal to or lower than the median for all of the activating sites in a given cancer type. Pearson correlations were also calculated between activity scores calculated using the RoKAI z-score with targets from the combination of PhosphoSitePlus and NetworKIN and kinase activating sites or kinase protein levels. To assess differences between the correlations of the activity scores with activating sites and their correlations with kinase protein levels, paired (by kinase) two-tailed Wilcoxon rank sum tests were used.

### Correlation of kinase metrics to kinase inhibitor response in cell lines

Proteomics and phosphoproteomics datasets for the NCI60 cells lines were obtained from Supplementary Tables 3 and 2, respectively, from Frejno, et al.^64^. Phosphosites were aggregated by the combination of HGNC symbols with the 11mer sequence flanking the site by keeping the rows with the least number of missing values for the same gene symbol-11mer combination. In cases where there was a tie, the rows were averaged. The protein dataset was processed similarly except that the data was aggregated by the HGNC symbol alone. Sites and proteins with measurements in less than 20 cell lines were removed, and the phosphosite data was centered by the median value for each site. The site data was then used as input for kinase activity inference calculations as described above (*Computational methods for kinase activity inference*) using either the targets from PhosphoSitePlus alone or in combination with NetworKIN. Drug inhibitor response data from the GDSC was downloaded from the Sanger Institute website (https://www.cancerrxgene.org/downloads/drug_data)^63^ and filtered to NCI60 cell lines and to inhibitors that target kinases. Spearman rank correlations between AUROC values for inhibitor response and either kinase activity scores or protein levels were calculated for kinases having both measurements that are established targets of a given inhibitor.

## Data Availability

The data for the benchmark can be accessed within benchmarKIN (https://github.com/saezlab/benchmarKIN) and is available at: https://zenodo.org/uploads/12566560.

For the kinase-substrate libraries, PhosphoSitePlus was obtained via https://www.phosphosite.org/staticDownloads#, PTMsigDB was obtained via https://proteomics.broadapps.org/ptmsigdb/, iKiP-DB was obtained via https://pubs.acs.org/doi/suppl/10.1021/acs.jproteome.2c00198/suppl_file/pr2c00198_si_007.zip and NetworKIN was obtained via http://netphorest.science/download/networkin_human_predictions_3.1.tsv.xz and is also available in the forementioned Zenodo repository.

## Code Availability

The code for the analysis presented in this manuscript is available at https://github.com/saezlab/kinase_benchmark. The benchmarKIN package is available at https://github.com/saezlab/benchmarKIN, along with detailed tutorials describing the benchmarking approaches presented here (https://benchmarkin.readthedocs.io).

## Acknowledgements

This work was supported by the German Federal Ministry of Education and Research, particularly the LiSyM Cancer research core (BMBF, Funding number: 031L0257B), the Innovation Campus Health + Life Science Alliance Heidelberg Mannheim, the Cancer Prevention & Research Institute of Texas (CPRIT) Awards RP220050, National Institutes of Health (NIH) grants from the National Cancer Institute (NCI) U24 CA271076, and funding from the McNair Medical Institute at The Robert and Janice McNair Foundation.

## Authors contributions

S.M.D. performed the kinase activity inference and the perturbation-based benchmark approach. E.J.J. conceptualized and performed the tumor-based benchmark approach and the drug response study. K.P.M. performed the different site-to-protein normalization procedures with the support of K.K. and D.R.M. T.M.Y.-B. calculated the kinase library scores with the support of J.L.J. and L.C.C. M.G.-R. calculated the Phosformer scores. A.L. processed the Hijazi datasets and was, together with E.P., involved in discussions around the perturbation-based benchmark. S.R.S. curated the activating site list. W.J. analyzed the correlation of host proteins for common targets of the same kinase. J.T.L. provided support for the drug response study. A.D. provided feedback throughout the analysis. B.Z. and J.S.R. supervised the project. S.M.D. and E.J.J. wrote the manuscript, which has been revised by all authors.

