## Supplementary Figures for "Comprehensive evaluation of phosphoproteomic-based kinase activity inference"

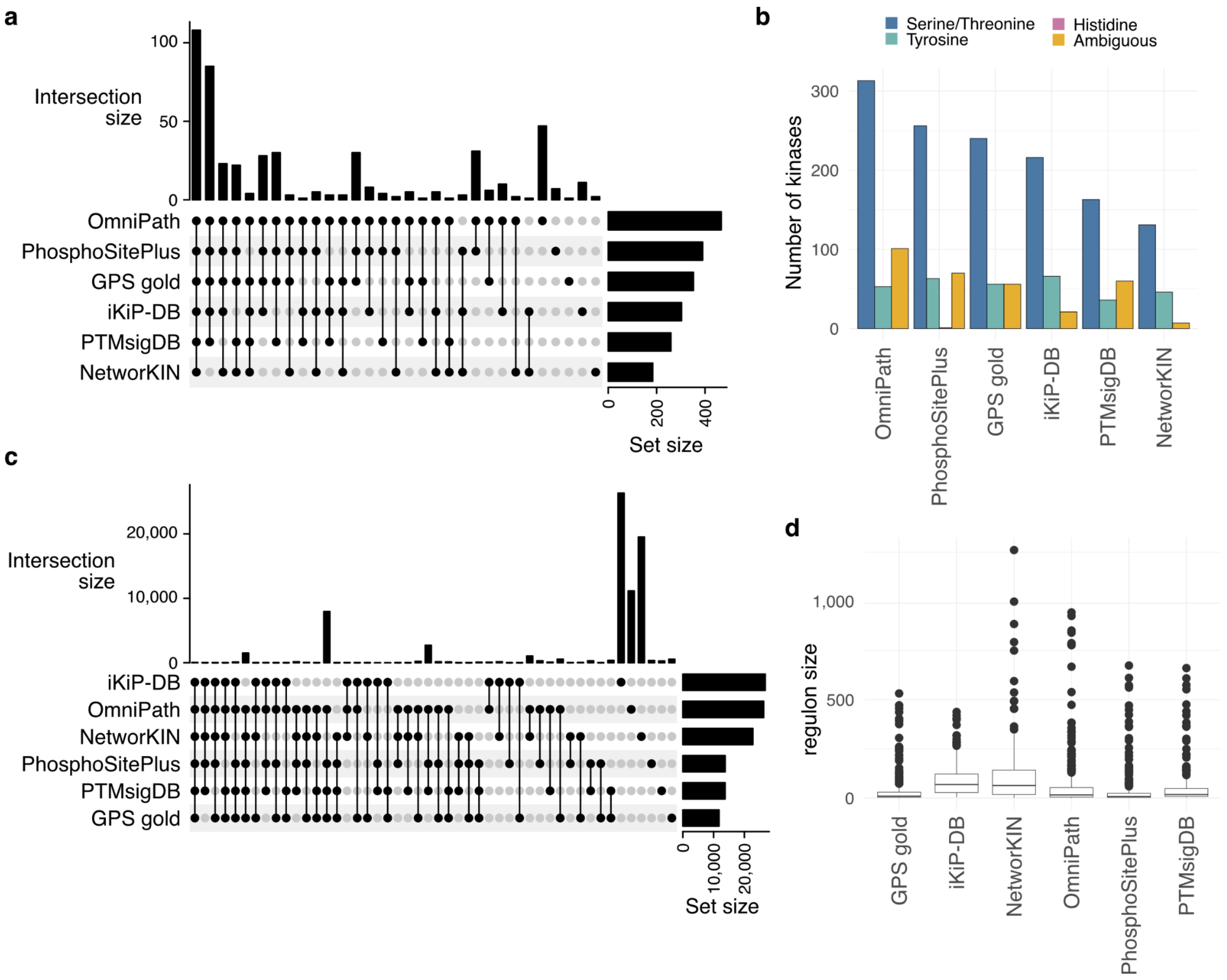


**Supplementary Figure 1  Kinase coverage and regulon size across resources.**

***a*** *UpSet plot visualizing the kinase intersections across kinase-substrate libraries.* ***b*** *Number of kinases covered in each resource, divided according to the different kinase classes (Serine/threonine, histidine, tyrosine or ambiguous).* ***c*** *UpSet plot visualizing the kinase-substrate interaction intersections across kinase-substrate libraries.* ***d*** *Regulon size, denoting the number of downstream phosphorylation sites, attributed to each kinase in every resource.*

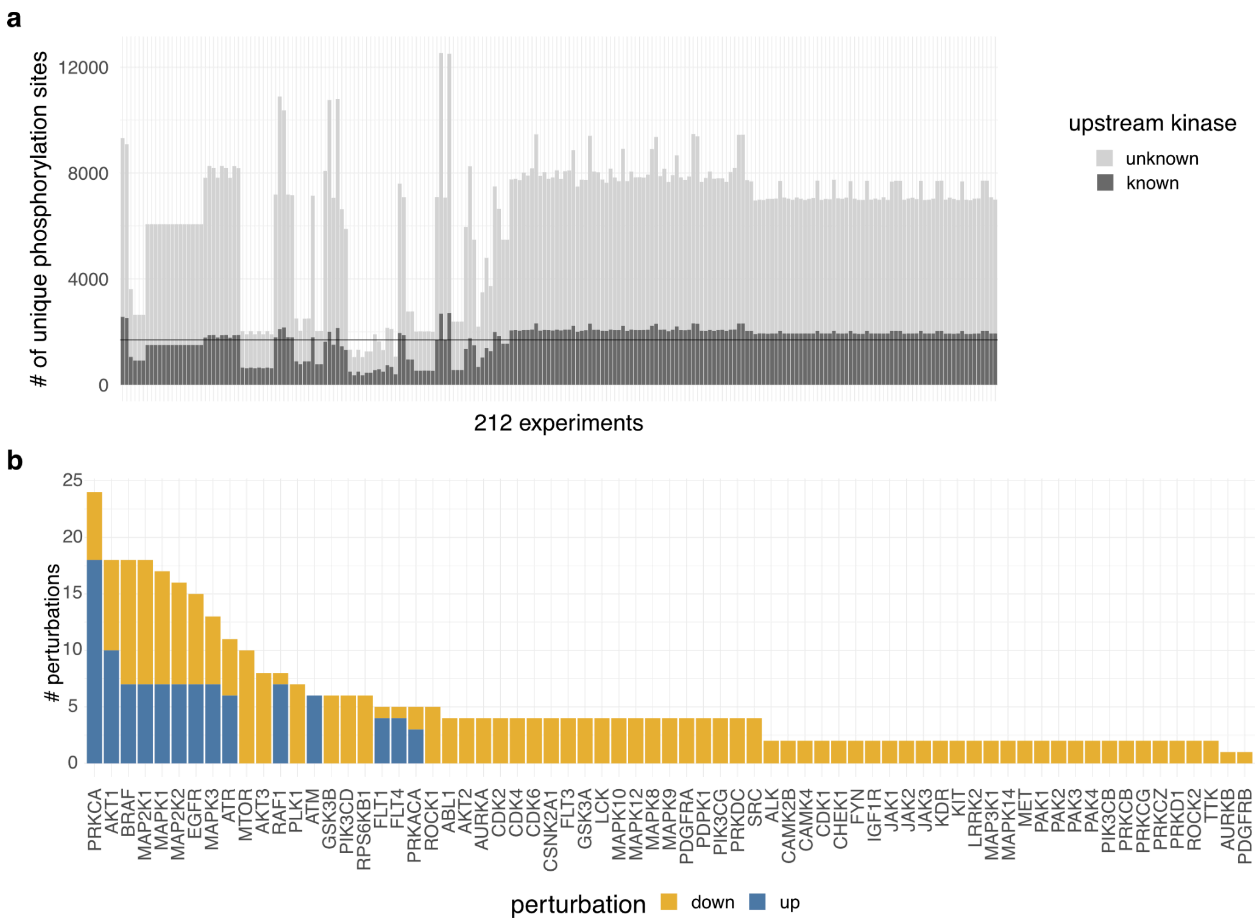


**Supplementary Figure 2 Overview perturbation datasets.**

***a*** *Number of phosphorylation sites identified in each experiment collected by Hernandez-Armenta et al. and Hijazi et al. Number of phosphorylation sites with a known upstream kinase in any of the kinase-substrate libraries (PhosphoSitePlus, PTMsigDB, GPS gold, OmniPath, iKiP-DB, NetworKIN) are highlighted in dark gray.* ***b*** *Number of perturbations for each kinase across all perturbations experiments of Hernandet-Armenta et al. and Hijazi et al. Perturbations associated with an increase in activity (up-regulation) are colored in blue and perturbations associated with a decrease in activity (down-regulation) are colored in orange.*

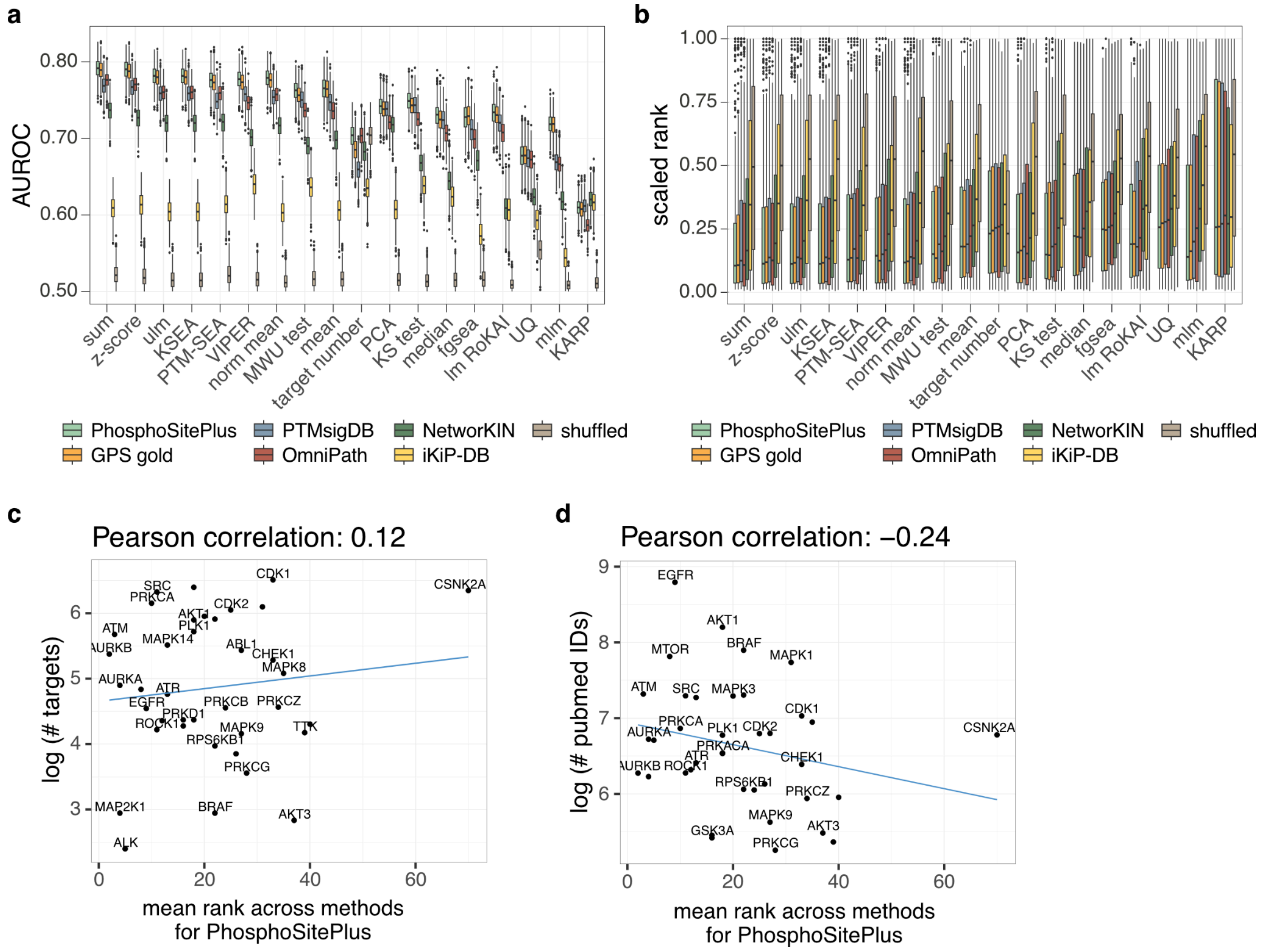


**Supplementary Figure 3 Evaluation of kinase activity inference.**

***a*** *Predictive performance of methods for kinase activity inference in identifying perturbed kinases from phosphoproteomics data. AUROC for each kinase-substrate library - computational algorithm prediction.* ***b*** *Scaled rank for each kinase-substrate library - computational algorithm prediction. The scaled rank is determined by taking the rank of the perturbed kinase in its experiment based on the activity, dividing it by the total number of kinases for which an activity was calculated.* ***c*** *Effect of number of targets on kinase rank. Correlation between the mean rank of each perturbed kinase across experiments and methods for PhosphoSitePlus and the number of targets in PhosphoSitePlus for that kinase.* ***d*** *Effect of study bias on kinase rank. Correlation between the mean rank of each perturbed kinase across experiments and methods for PhosphoSitePlus and the number of studies related to that kinase.*

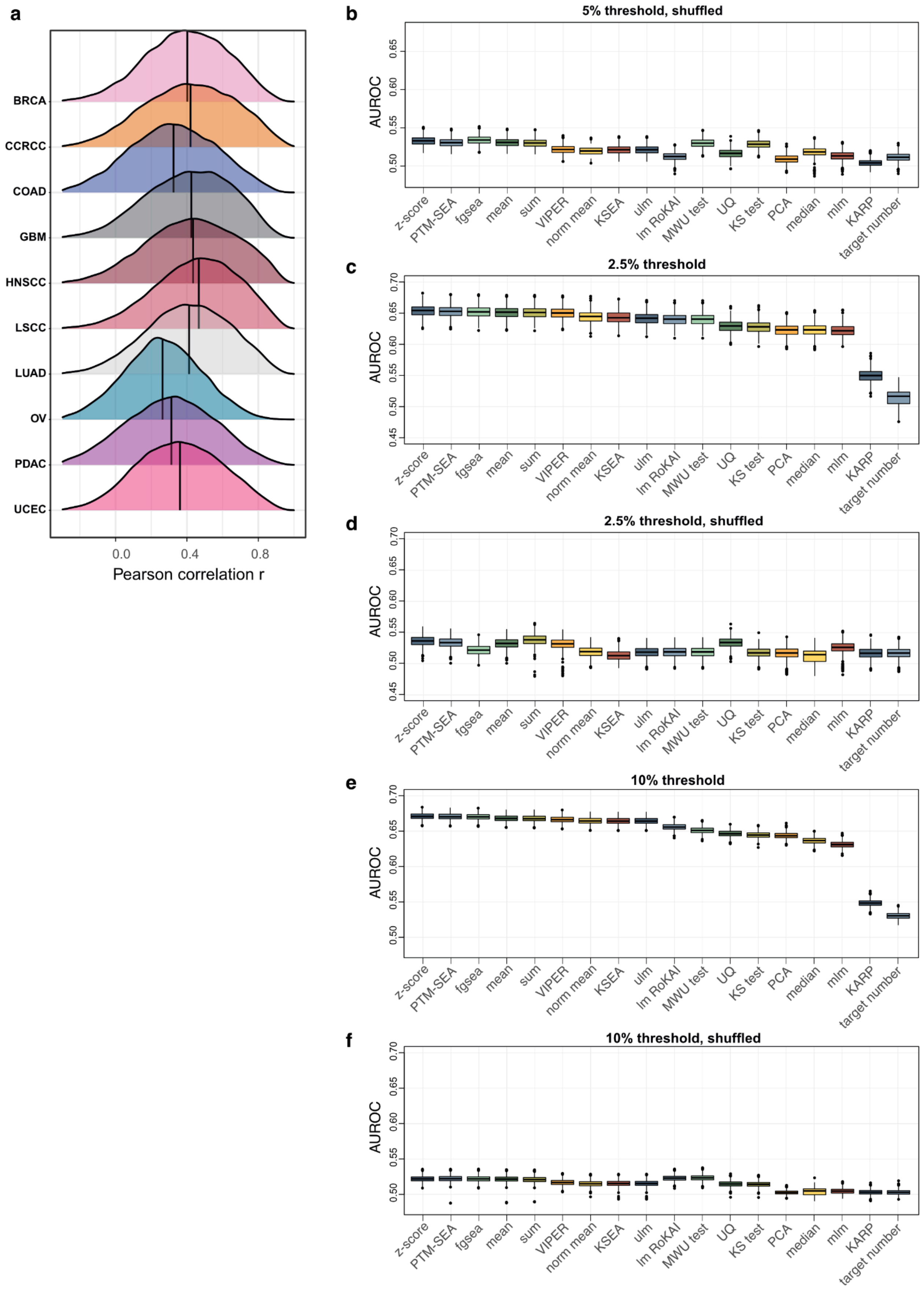
a
BRCA
CCRCC
COAD
GBM
HNSCC
LSCC
LUAD
OV
PDAC
UCEC
0.0
0.4
0.8
Pearson correlation r
b
5% threshold, shuffled
c
2.5% threshold
d
2.5% threshold, shuffled
e
10% threshold
f
10% threshold, shuffled
AUROC
z-score
PTM-SEA
fgsea
mean
sum
VIPER
norm mean
KSEA
ulm
lm RoKAI
MWU test
UQ
KS test
PCA
median
mlm
KARP
target number

**Supplementary Figure 4  Testing tumor-based benchmark approach**

**a** Distributions of correlations between phosphosites and host proteins for each of the ten cancer types profiled by CPTAC. **b** Sample AUROCs for scores calculated using shuffled targets from PhosphositePlus. **c** Sample AUROCs for an alternative gold standard set established using the top and bottom 2.5% of protein levels. **d** Sample AUROCs for scores calculated using shuffled targets from Phosphosite Plus and the 2.5% threshold for the gold standard. **e** Sample AUROCs for an alternative gold standard set established using the top and bottom 10% of protein levels. **f** Sample AUROCs for scores calculated using shuffled targets from PhosphositePlus and the 10% threshold for the gold standard.

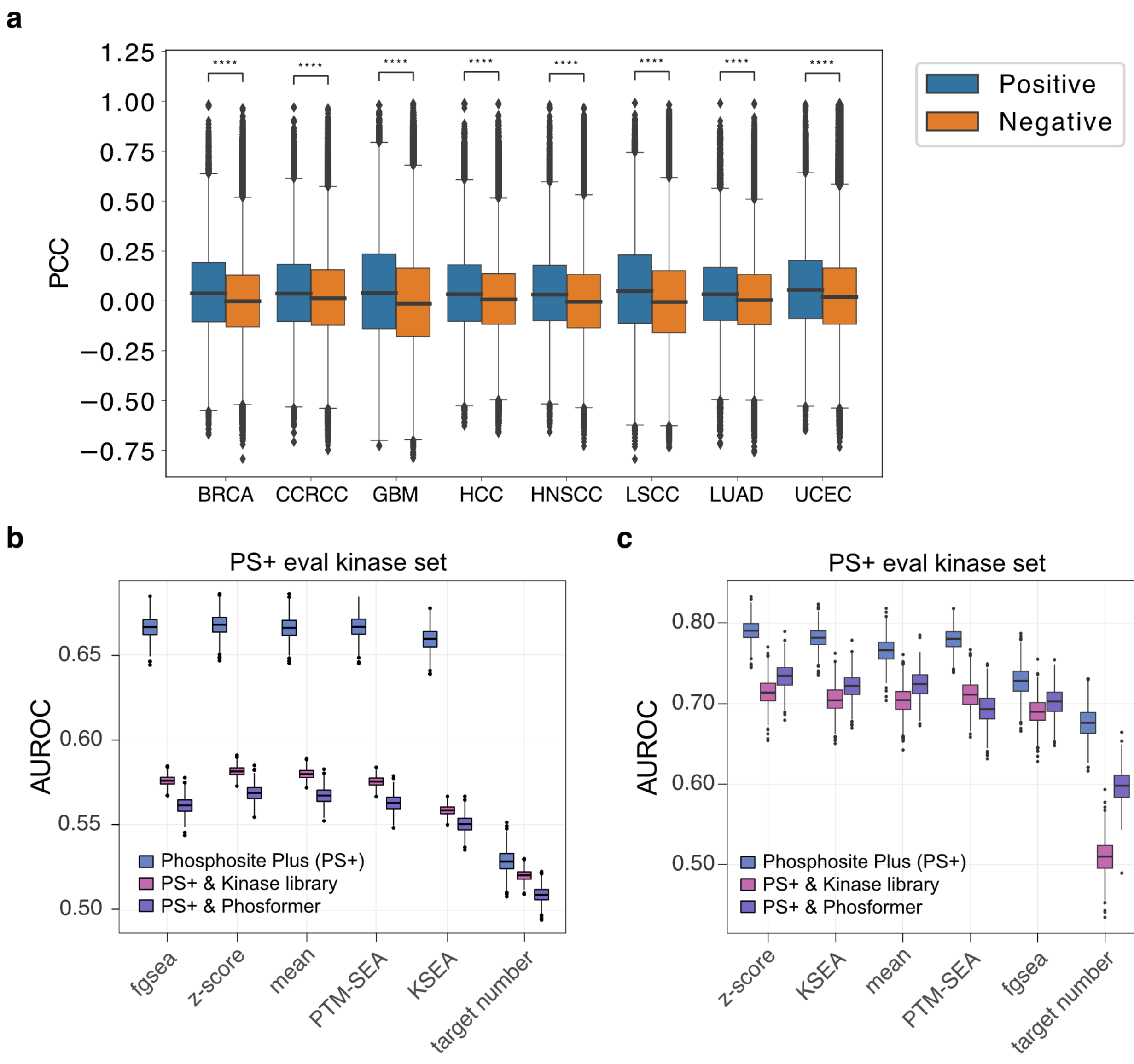


**Supplementary Figure 5 Protein correlation and evaluation of predicted phosphorylation sites.** ***a*** *Pearson correlations for pairs of proteins that contain sites that are common targets of the same kinase (blue) vs. correlations for pairs of proteins that contain targets from different kinase groups.* ***b*** *AUROCs for various methods for kinase activity inference using PhosphositePlus in combination with the Kinase Library or Phosformer in the tumor-based benchmark approach.* ***c*** *AUROCs for various methods for kinase activity inference using PhosphositePlus in combination with the Kinase Library or Phosformer in the perturbation-based benchmark approach.*

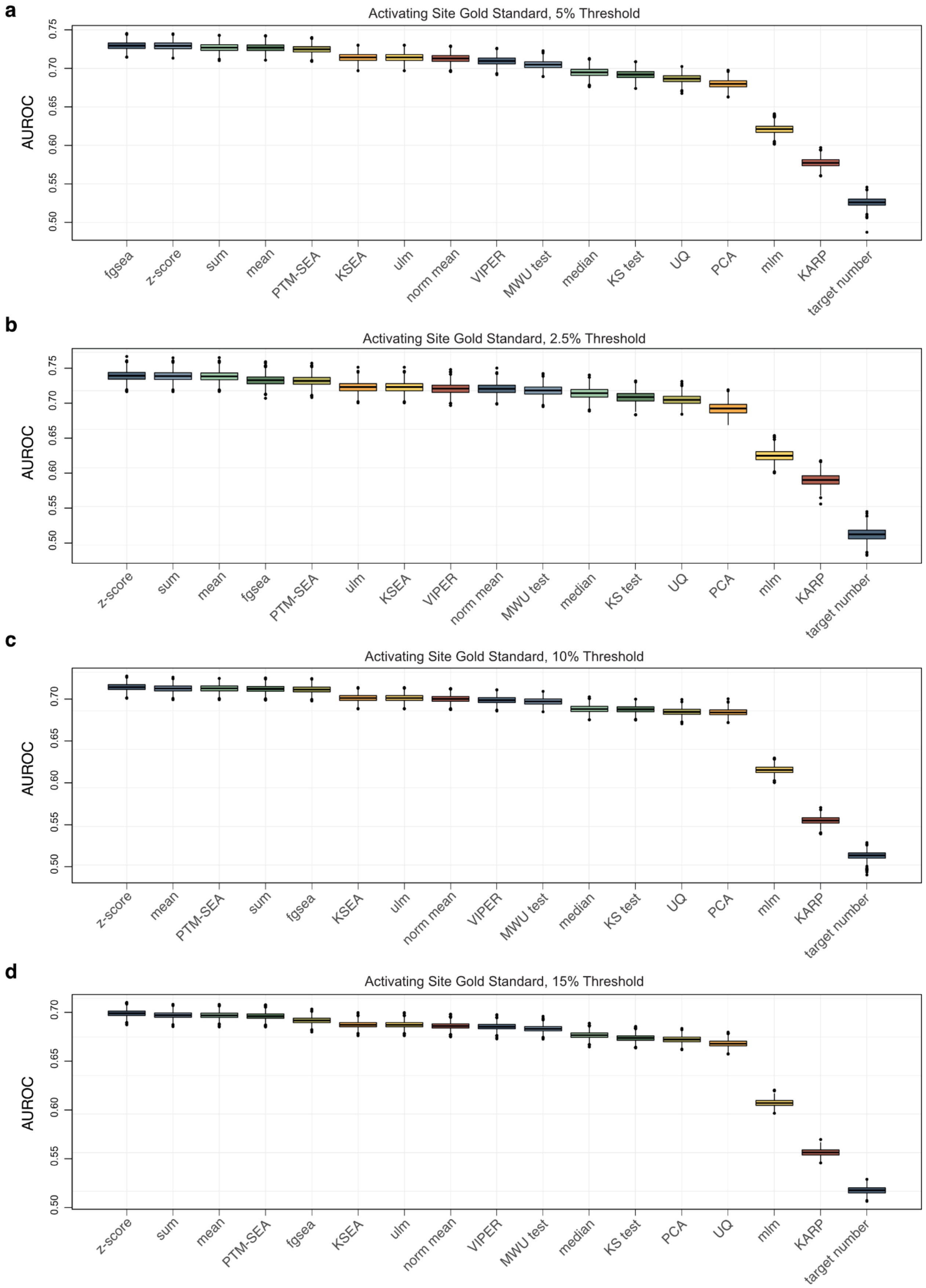
a
Activating Site Gold Standard, 5% Threshold
AUROC
0.50
0.55
0.60
0.65
0.70
0.75
fgsea
z-score
sum
mean
PTM-SEA
KSEA
ulm
norm mean
VIPER
MWU test
median
KS test
UQ
PCA
mlm
KARP
target number
b
Activating Site Gold Standard, 2.5% Threshold
AUROC
0.50
0.55
0.60
0.65
0.70
0.75
z-score
sum
mean
fgsea
PTM-SEA
ulm
KSEA
VIPER
norm mean
MWU test
median
KS test
UQ
PCA
mlm
KARP
target number
c
Activating Site Gold Standard, 10% Threshold
AUROC
0.50
0.55
0.60
0.65
0.70
z-score
mean
PTM-SEA
sum
fgsea
KSEA
ulm
norm mean
VIPER
MWU test
median
KS test
UQ
PCA
mlm
KARP
target number
d
Activating Site Gold Standard, 15% Threshold
AUROC
0.50
0.55
0.60
0.65
0.70
z-score
sum
mean
PTM-SEA
fgsea
KSEA
ulm
norm mean
VIPER
MWU test
median
KS test
PCA
UQ
mlm
KARP
target number

**Supplementary Figure 6  Activating sites thresholds**

Sample AUROCs for scores calculated using the combination of PhosphositePlus and NetworKIN targets and evaluated using an alternative gold standard set established using the top and bottom 5% **a**, 2.5% **b**, 10% **c**, or 15% **d** of activating sites on kinases instead of kinase protein levels.
